# Energy scarcity and impaired mitochondrial translation induce perinuclear stress granule clustering

**DOI:** 10.1101/2024.02.01.578399

**Authors:** Uxoa Fernandez-Pelayo, Mikel Muñoz-Oreja, Marina Villar-Fernandez, Amaia Lopez de Arbina, Irati Aiestaran-Zelaia, María Jesús Sánchez-Guisado, Boris Pantic, Amaia Elicegui, Monica Zufiria, Pablo Iruzubieta, Maialen Sagartzazu-Aizpurua, Jesús M. Aizpurua, Matthew Gegg, Sonia Alonso-Martin, Jesus Ruiz-Cabello, Francisco Gil-Bea, Antonella Spinazzola, Adolfo Lopez de Munain, Ian James Holt

## Abstract

Many proteins linked to amyotrophic lateral sclerosis and fronto-temporal dementia (ALS-FTD) change their cellular location and coalesce in cytoplasmic inclusion bodies in the disease state; yet the factors that govern protein relocation and organization remain unclear. Here, we show that inhibition of glycolysis and mitochondrial protein synthesis causes many proteins involved in ALS-FTD to change location, and form a novel structure comprising a ring of stress granules encircling the aggresome, a focal microtubule-based structure beside the nucleus. A perinuclear ring of stress granules also forms in activated microglia of mice exposed to the glycolytic inhibitor, 2-Deoxy-D-glucose. We propose that the new arrangement increases the risk of the stress granules merging and converting from the liquid phase to the insoluble inclusion characteristic of ALS-FTD. Thus, our findings suggest that that compromised nutrient and energy metabolism can precipitate a molecular cascade that ultimately leads to the pathological hallmark of ALS-FTD the perinuclear inclusion body.

**Graphical abstract:** 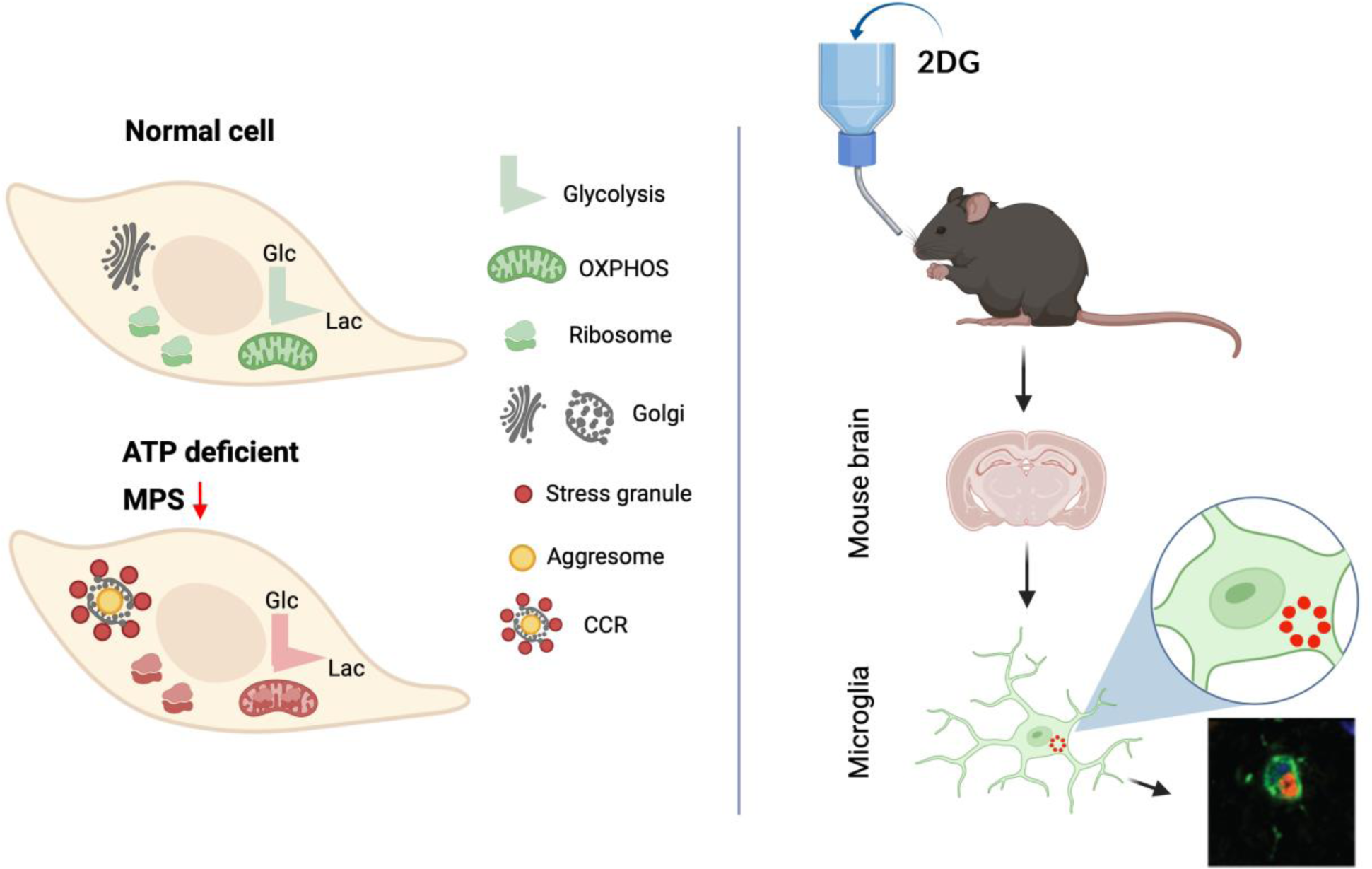

Inhibition of glycolysis and mitochondrial protein synthesis induces translocation of a swathe of ALS-FTD related proteins in primary human fibroblasts. The relocated proteins form concentric cytoplasmic rings (CCR) comprising stress granules, the Golgi and the aggresome, beside the nucleus. A perinuclear ring of stress granules forms in the mouse brain following intermittent nutrient restriction, with the glucose analog 2DG. The CCR is potentially a key intermediate step in the formation of pathological inclusions and so perturbed nutrient and energy metabolism encompassing impaired mitochondrial translation could precipitate the ALS-FTD disease cascade.

## INTRODUCTION

Amyotrophic lateral sclerosis (ALS) is a devastating and incurable neurodegenerative disorder, resulting from motor neuron degeneration. ALS overlaps clinically and genetically with frontotemporal dementia (FTD), and together they form part of a wider group of neurodegenerative disorders known as proteinopathies (Ross and Poirier, 2004). The major pathological hallmark of ALS is cytoplasmic plaques of ubiquitinated proteins that in a majority of affected individuals contain nuclear RNA binding proteins, principally TDP-43 (TAR DNA binding protein 43) (Ling et al., 2013). FUS and hnRNPA1, that are mutated in some familial forms of ALS, that account for around 5% and less than 1% of familial cases, respectively (Andersen and Al-Chalabi, 2011), are also frequent components of the pathological inclusions (Aulas and Vande Velde, 2015; Blokhuis et al., 2013; Peters et al., 2015). These nuclear proteins are among those that relocate to the cytoplasm where they form condensates with RNA (stress granules) upon the inhibition of protein synthesis, in response to stresses such as Sodium arsenite and heat (e.g. (Mateju et al., 2017)). Stress granule induction is linked to mTOR signaling (Wippich et al., 2013), and the HSPB8-BAG3-HSP70 Chaperone Complex maintains their dynamism (Ganassi et al., 2016), allowing them to be disassembled rapidly by activated Valosin-containing protein (VCP/p97) (Wang et al., 2019a).

Although stress granule proteins in the cytoplasm, such as TDP-43, are ordinarily highly mobile, and readily reversible, when mutated or misfolded they can become static making them more difficult to disassemble; and such aberrant stress granules can then only be cleared by the protein quality control systems (e.g. (Chitiprolu et al., 2018; Ganassi et al., 2016; Mateju et al., 2017; Turakhiya et al., 2018; Wang et al., 2019a). Hence, it has been proposed that ALS cytoplasmic inclusions are derived from aberrant stress granules that the cell failed to re-cycle (Li et al., 2013; Vanderweyde et al., 2013). Additionally, other familial forms of ALS are linked to defective stress granule clearance such as Optineurin, Ataxin 2, SQSTM1 (Ubiquitin binding protein sequestosome 1), Ubiquilin-2, VAPB (VAMP associated protein B/C), or VCP (Johnson et al., 2010; Ju et al., 2008). Notwithstanding all the above, the role of stress granules in the formation of TDP-43 pathological inclusions remains a matter of debate (Gasset-Rosa et al., 2019; Zhang et al., 2019).

Several of the proteins listed above are associated with a stress-induced structure, formed of microtubules, called the aggresome (Kopito, 2000), which segregates misfolded, ubiquitinated proteins from the rest of the cell, and recycles them enhancing cell survival (Kawaguchi et al., 2003; Olzmann et al., 2008). In cancer cells, the aggresome can be induced by proteasome inhibition with MG132 (Johnston et al., 1998; Taylor et al., 2003). Stress granules and the aggresome rarely form in the same cells, although a handful of studies suggest that stress granules can be trafficked along microtubules for recycling at the aggresome (Kawaguchi et al., 2003; Kwon et al., 2007; Mateju et al., 2017). On the other hand, while stress granules and inclusions of mutant ubiquitin form after heat shock or MG132 treatment in some cell types, they are spatially and temporally distinct, and so their relationship to one another is unclear (Mateju et al., 2017).

Although the natural triggers of the cytoplasmic inclusions seen in ALS are not known, one potential contributor is impaired energy metabolism, as perturbed nutrient metabolism, altered mitochondrial transport and morphology, have all been reported in ALS patients or ALS models (Hor et al., 2021; Magrane et al., 2014; Palamiuc et al., 2015; Watanabe et al., 2016). Moreover, inhibition of glycolytic or mitochondrial energy production has been used to induce stress granules in human cancer cells and rodent cells (Kedersha et al., 2002; Wang et al., 2019b; Wang et al., 2022), and AMPK activation, which is a sensor of ATP depletion, has been implicated in stress granule formation (Kuo et al., 2020; Mahboubi et al., 2016). On the other hand, stress granule formation is an ATP-dependent process (Jain et al., 2016).

Here, we restricted energy production in primary human fibroblasts to determine the effect on the location and organization of a suite of ALS-related proteins, including those associated with stress granules. We show that in primary human fibroblasts inhibition of glycolysis and mitochondrial protein synthesis induces a ring of stress granules to form around a microtubule-based structure, the aggresome, adjacent to the nucleus. In the murine brain, the glycolytic inhibitor 2-Deoxy-D-glucose (2DG) induces the formation of a similar perinuclear ring of stress granules, or a single body similar in appearance to those seen in ALS-FTD in affected cells. These findings indicate how stress granules can cluster close to the nucleus, which is potentially a key intermediate step in the formation of pathological inclusions. As a corollary, the results suggest that impaired energy metabolism could precipitate a cascade of molecular events that leads to ALS-FTD-related inclusions.

## RESULTS

### Inhibition of glycolysis and mitochondrial respiration induces relocation of TDP-43 and FUS to the cytoplasm

To determine the effect of energy metabolism on the distribution of proteins involved in ALS, we targeted, individually or together, the two cellular pathways, glycolysis and respiration, that produce ATP. Glycolysis, among other pathways, is inhibited by 2DG, and 2DG is sufficient to induce stress granules in rodent renal tubular cells (Wang et al., 2019b). However, while 2DG treatment of primary human fibroblasts increased SQSTM1 expression and extended its distribution to the nucleus, it did not affect the cellular location of the RNA binding proteins TDP-43 or FUS, nor did stress granules containing the canonical marker G3BP1 form, when fibroblasts were exposed to 2DG at a concentration of 20 mM for 24 hours or 200 mM for 3 hours (Fig. S1A-C). TDP-43 or FUS also remained in the nucleus when we depolarized the mitochondria by treating the cells with 10 μM of the proton ionophore CCCP (Carbonyl cyanide m-chlorophenyl hydrazone) (Fig. S1D). Instead, combining 200 mM 2DG and 10 μM CCCP caused almost all the TDP-43 and FUS to disperse throughout the cytoplasm without inducing SG formation (Fig. S1E, S1F), concordant with an earlier report that found that ATP supports the nuclear localization of TDP-43 and FUS and that ATP is necessary for stress granule formation (Jain et al., 2016).

CCCP promotes autophagic clearance of mitochondria (Narendra et al., 2008) and is thermogenic, as well as completely abrogating mitochondrial ATP production; therefore, in a more targeted and less extreme approach, we restricted mitochondrial ATP production via respiratory chain complex I with Rotenone. Three-hour treatments of 200 mM 2DG with 0.1 μM Rotenone also caused the nuclear RNA binding protein FUS to disperse throughout the cytoplasm in the majority of cells (Figs. 1a, S2A). However, when fibroblasts were exposed to a ten-fold lower dose of 2DG (20 mM) for 24 hours and 0.1 μM Rotenone throughout this period, or for the final 2-3 hours, TDP-43 and FUS formed cytoplasmic foci coincident with G3BP1; i.e., stress granules (Figs. 1a, S2B). The stress granules were generally spaced well apart, towards the cell periphery (Fig. S2B). Because 2DG has pleiotropic effects beyond inhibition of glycolysis (Pantic et al., 2021), we also tested a more specific inhibitor of glycolysis, Koningic acid (KA) (Endo et al., 1985). Treatments of 0.1 μM Rotenone in combination with 1 μM KA for 24 hours, induced stress granules in control fibroblasts, similar to 2DG (Fig. S2C). The minimum time to induce stress granules with 20 mM 2DG and 0.1 μM Rotenone was 90 minutes, while none were seen with 2DG alone at any time point (Fig S2D). Hence, co-inhibition of glycolysis (20 mM 2DG or 1 μM KA) and respiratory complex I (0.1 μM Rotenone) induces stress granules in primary human fibroblasts, whereas harsher treatments with 200 mM 2DG and CCCP or Rotenone result in TDP-43 and FUS dispersal in the cytoplasm without stress granule formation (Figs. 1a, S2E, S2F).

**Figure 1.**
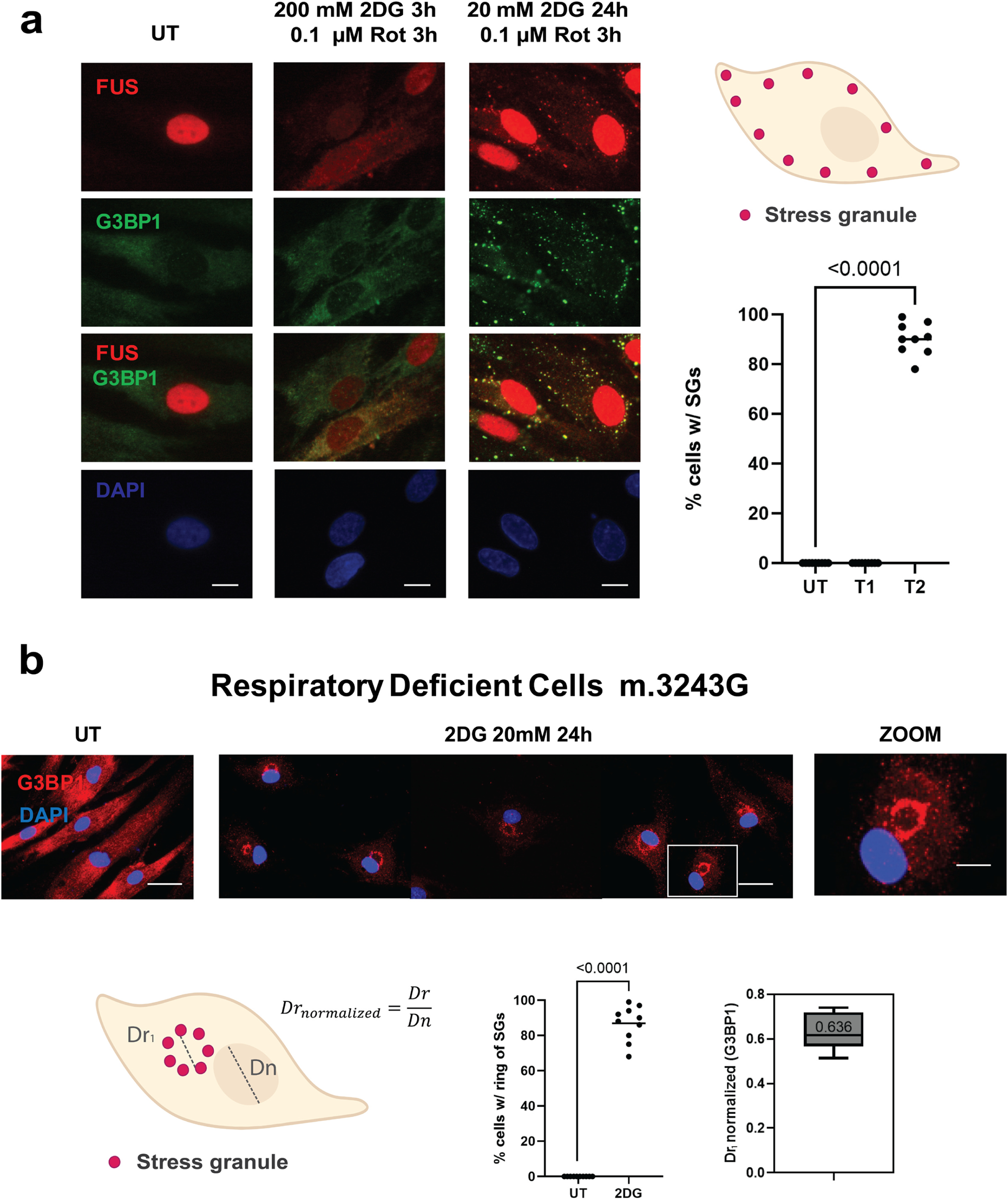
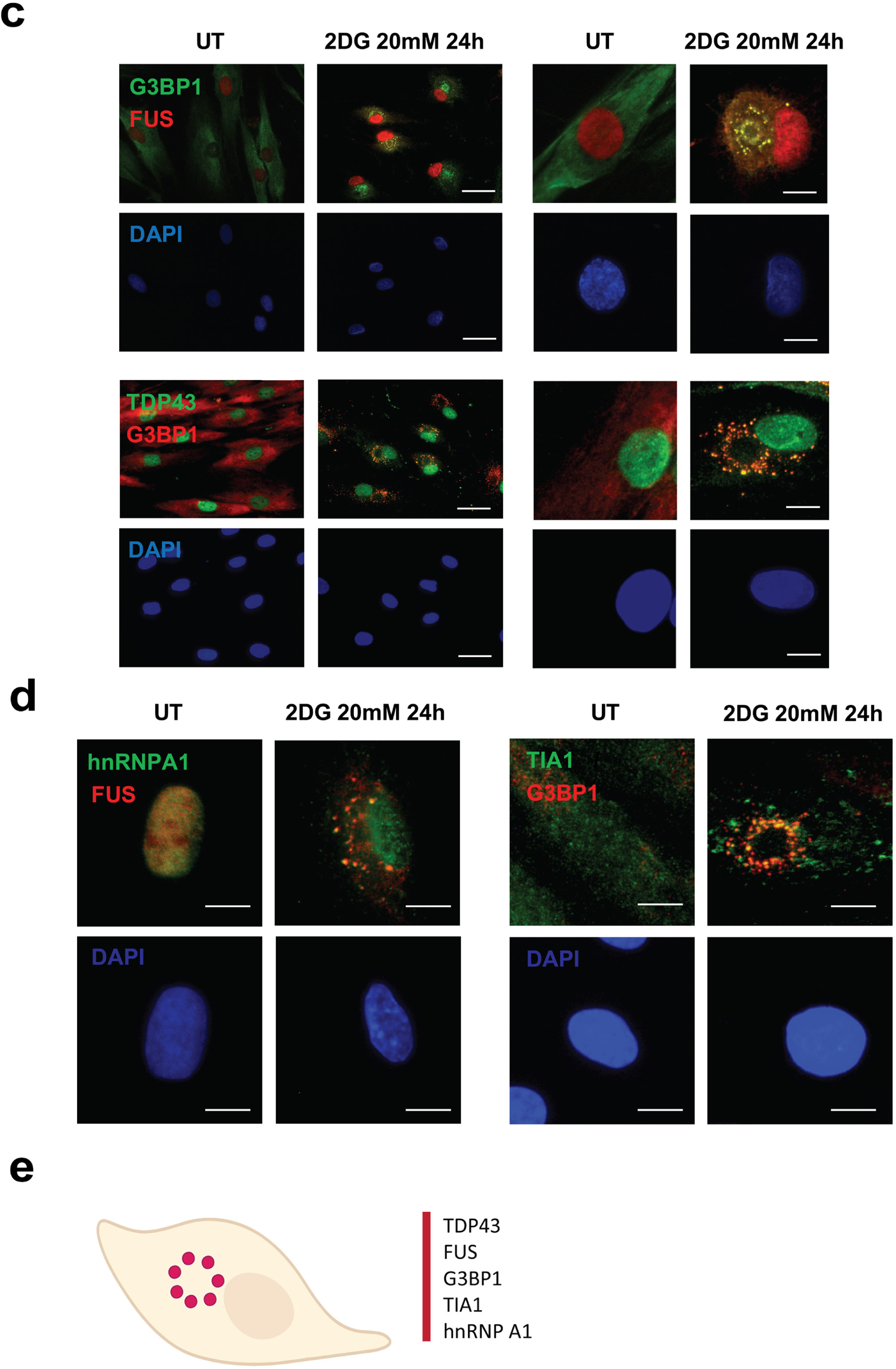
Restricted energy metabolism induces widely dispersed stress granules in control fibroblasts, or a perinuclear ring in respiratory deficient cells owing to a genetic defect in the mitochondrial DNA. **a**) Control fibroblasts were left untreated (UT), or exposed to 0.1 μM Rotenone with 200 mM 2DG for 3h (T1), or to 0.1 μM Rotenone for the final 3 h of a 24 h, 20 mM 2DG treatment (T2). Fixed cells were immunostained for the canonical stress granule marker G3BP1 (green) and the nuclear RNA binding protein FUS (red), and mounted with DAPI to mark the nuclei. Scale bar 30 µm. Chart to the right indicates the proportion of cells with (w) stress granules in the case of T2 (measured in 450 cells across n=9 independent experiments in 6 control cell lines). **b**) Respiratory deficient human fibroblasts (m.3243G) were treated with 20 mM 2DG for 24 h, after fixation the cells were stained an antibody to G3BP1 (red) and DAPI (blue), and the former revealed a perinuclear ring of stress granules in 87% of the cells. Analysis of 500 cells, across n=10 independent experiments on 3 mutant cell lines. The diameter to the outer edge of the ring was measured and was normalized to the diameter of the nucleus in each cell, as represented in the box chart. Analysis of G3BP1 signal in 300 cells, across n=10 independent experiments on 2 mutant cell lines. To the left of the charts, an illustration of the stress granule ring with the formula for calculating its diameter (*Dr*). **c**) Representative images of 180 cells treated as panel **b**, immunostained for TDP-43 or FUS, and G3BP1, as indicated. n=6 independent experiments on 2 mutant cell lines for each immunostaining. **d**) As panel **c**, immunostained for TIA1 and G3BP1, or hnRNPA1 and FUS. 150 cells, n=5 independent experiments on 2 mutant cell lines for each immunostaining. Scale bars in panels b and c: 50 µm and 30 µm in zoomed images to the right, and 30 µm in panel d. **e**) Illustration of the identified components of the cytoplasmic rings.

### Inhibition of glycolysis induces the formation of concentric perinuclear rings containing ALS associated factors in cells with a genetic defect in mitochondrial protein synthesis

As a genetic model of complex I deficiency, we analyzed respiratory deficient fibroblasts carrying a pathological mitochondrial DNA point mutant (m.3243G) (Pantic et al., 2021). In the m.3243G fibroblasts, 2DG (without Rotenone) induced cytoplasmic stress granules (Fig. 1b). 1 µM KA for 24 h proved highly toxic to respiratory deficient fibroblasts; nevertheless, FUS and G3BP1 were concentrated close to the nucleus in the surviving cells, similar to the effect of 2DG (Fig. S3). Typically, the stress granules formed a perinuclear cluster or ring in m.3243G cells, a distribution that was not seen in the control fibroblasts exposed to 2DG with Rotenone (Fig. 1b, S3 versus Figs. 1a, S2B-S2D). Defining a perfect circle as 1, the circularity of the perinuclear ring of stress granules is 0.69, with a diameter of 14±3 µm, two-thirds that of the nucleus, and the distance between the nuclear envelope and the ring of stress granules ranged from 0-4.6 µm (Fig. 1b). The thickness of each ring is around a quarter of the outer diameter of each ring, with a maximum of a third of the outer diameter. Both FUS and TDP-43 were detected in the ring of stress granules, although the majority of each protein remained in the nucleus (Fig. 1c). TIA1 and hnRNPA1, two other nuclear RNA binding proteins linked to ALS that have been found in stress granules, also colocalized with G3BP1 in the rings, in the 2DG-treated respiratory deficient cells (Fig. 1d), further supporting the conclusion that these structures are archetypal stress granules, albeit with a highly unusual distribution (Fig 1e).

The perinuclear region of the cell, where the ring of stress granules formed, is the usual position of the Golgi, and immunostaining for the Golgi markers RCAS1 and GORASP2 indicated that the rings surrounded a reorganized Golgi apparatus, with a diameter approximately two-thirds that of the stress granule ring (Fig. 2a vs. Fig. 1b). Inhibition of energy production in control fibroblasts also resulted in Golgi reorganization and circularization (Fig. 2a, Fig S4A). Superoxide dismutase 1 (SOD1) was one of the first factors linked to familial ALS (Rosen, 1993), and its cellular distribution was similarly affected by energy scarcity. In response to restricted energy production, SOD1 colocalized with the Golgi markers inside the ring of stress granules (Fig. 2b and 2d).

**Figure 2.**
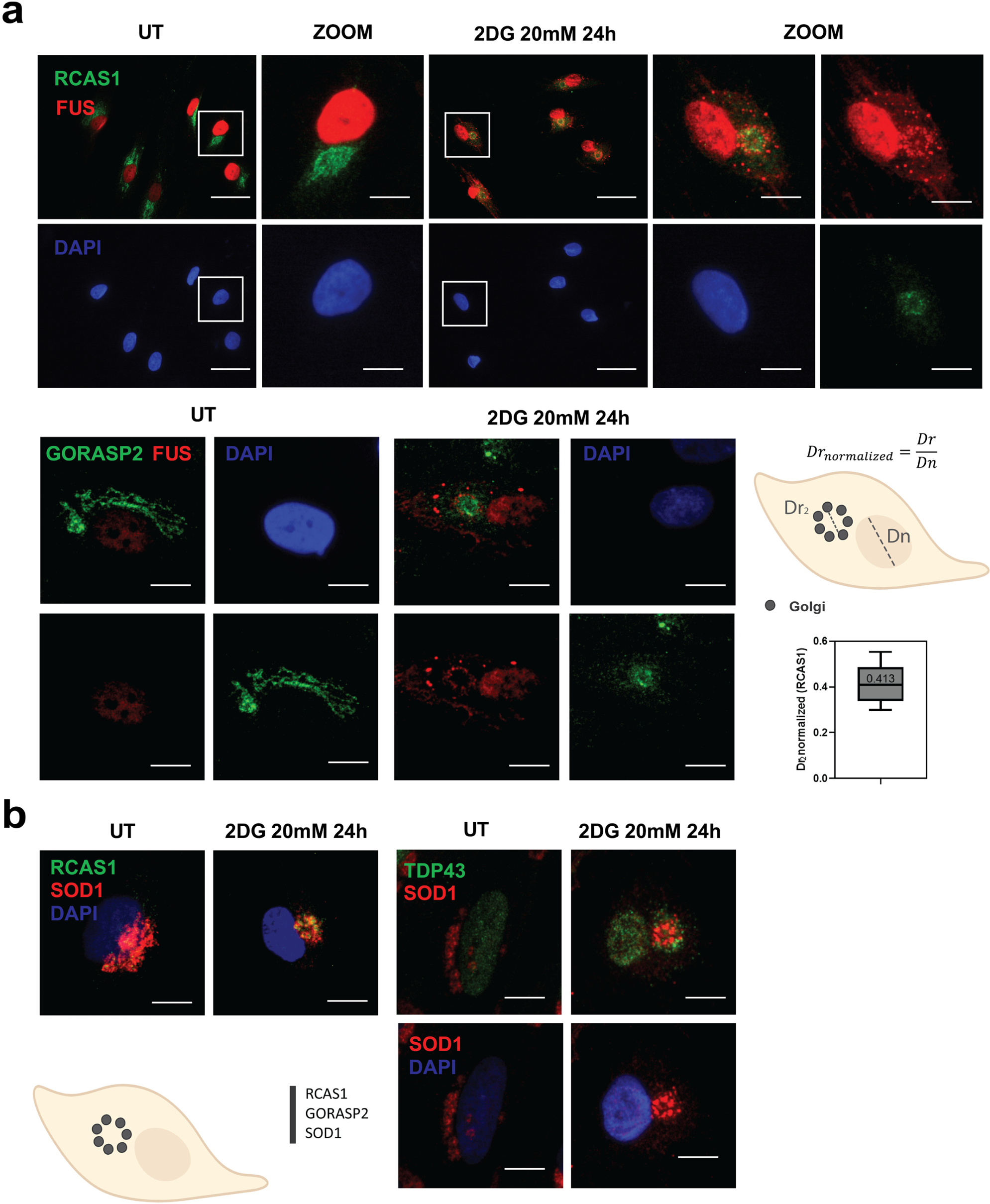
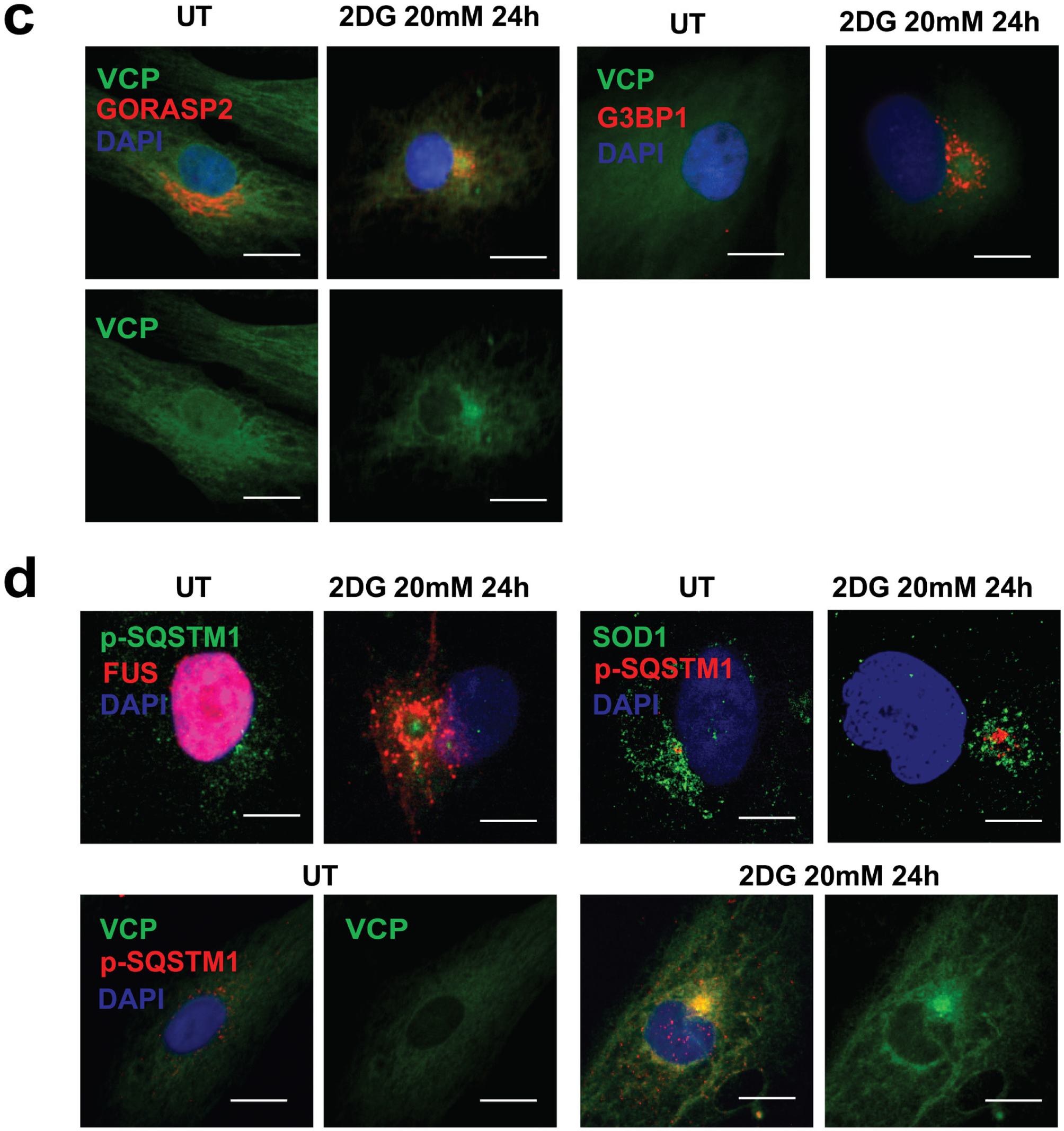
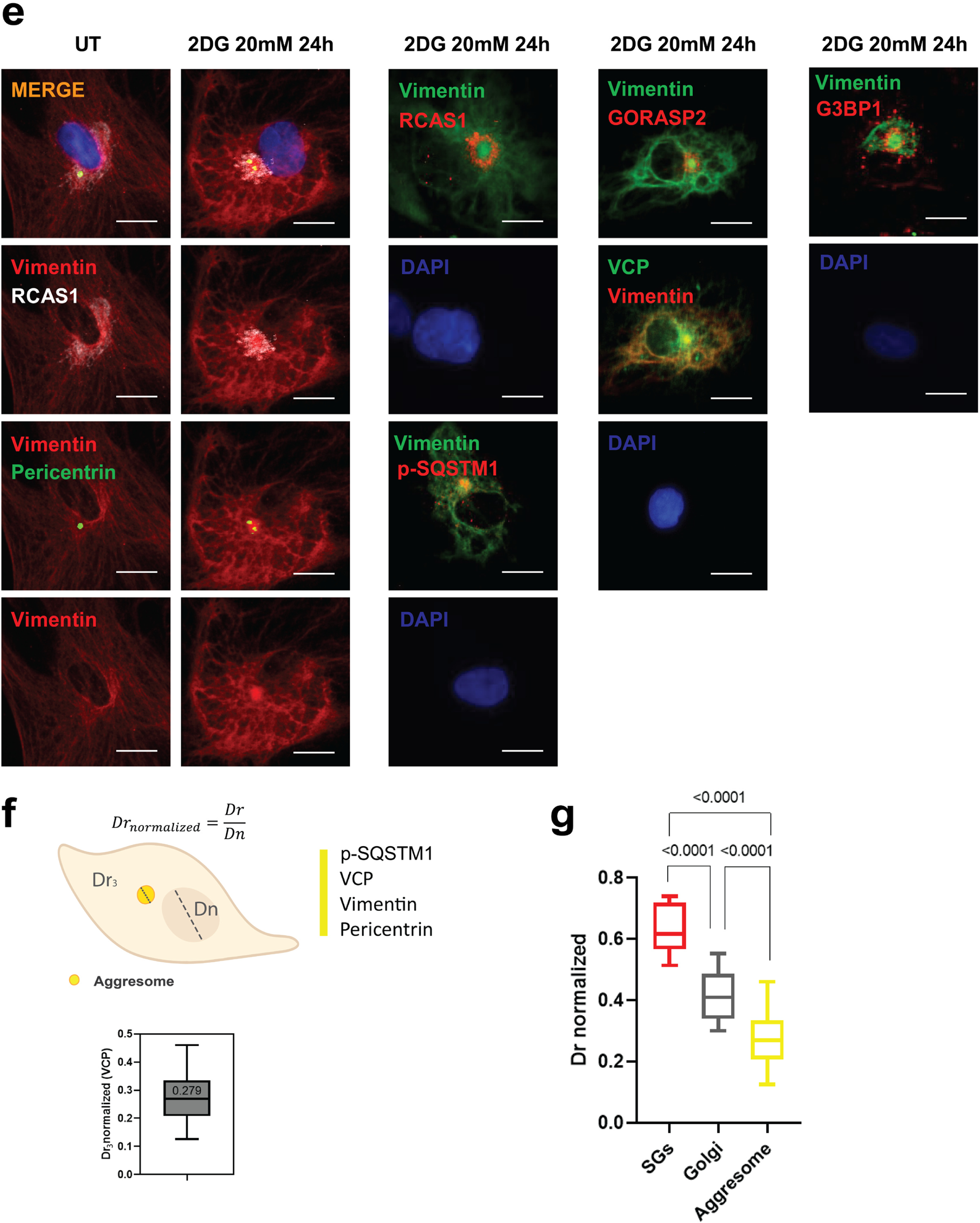
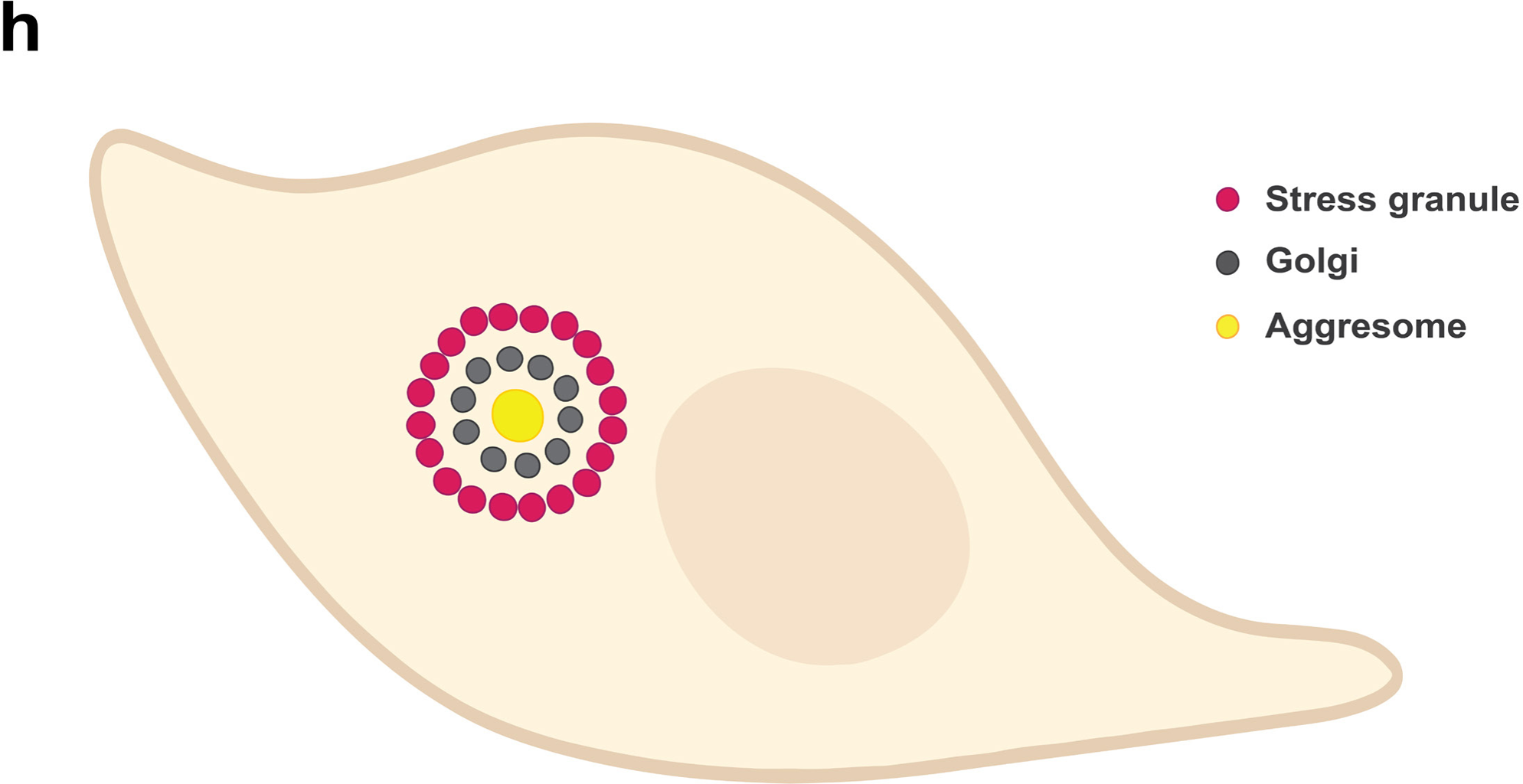
Restricted energy metabolism remodels the Golgi and induces the aggresome. Respiratory deficient (m3243G) human fibroblasts were treated with 20 mM 2DG for 24 h. After fixing, cell nuclei were stained blue with DAPI and immunostained for the Golgi markers, RCAS1 and GORASP2, and FUS, TDP-43, SOD1, VCP/p97, phosphorylated SQSTM1 (p-SQSTM1), Vimentin and the MTOC marker pericentrin, as indicated. Representative images from an analysis of a minimum of 180 cells, across n=6 independent experiments for each immunostaining, except in the case of p-SQSTM1 (100 cells, n=4 independent experiments) (**a-e**). Scale bars: 30 µm. The cell images of panel **a** are accompanied by a drawing of the Golgi ring and measurements of its diameter (*Dr*) normalized to the nucleus, based on an analysis of RCAS1 signal in 200 cells (n=10 independent experiments on 2 mutant cell lines). Similarly, panel **f**) shows an illustration of the aggresome diameter based on an analysis of VCP signal in 200 cells (n=10 independent experiments). **g**) Chart comparing the diameters of the stress granule, Golgi and aggresome rings. **h**) Schematic diagram representing the juxtaposition of proteins in the rings, based on the immunostainings in panels **a-e**, and Fig. 1.

We next considered whether the stress granules and Golgi rings were arranged around the aggresome, which is a stress-induced perinuclear structure that receives ubiquitinated and misfolded proteins for recycling, as well as in some circumstances stress granules (Mateju et al., 2017). The aggresome is bounded by a Vimentin cage and associated with VCP and phosphorylated SQSTM1 (pSQSTM1), with the mitotic organizing center (MTOC) at its core (Hirabayashi et al., 2001; Johnston et al., 1998; Shin, 1998). Immunostaining of the 2DG-treated respiratory deficient cells revealed the MTOC marker pericentrin at the center of the rings, accompanied by a dense Vimentin disc bounded by the Golgi (marked with RCAS1 and GORASP2); similarly, some VCP was concentrated inside the ring of stress granules and the Golgi, together with the entire cellular complement of pSQSTM1 (Fig. 2c-2e, S4C). The mean normalized diameter of the VCP-labeled aggresome was 0.28 (Fig. 2f), almost a third smaller than the Golgi ring (*p*<0.0001) and half that of the stress granule ring (*p*<0.0001) (Fig. 2g). In contrast to the phosphorylated SQSTM1 that formed a single condensed disc or sphere in the 2DG-treated respiratory deficient cells, the bulk of the SQSTM formed scattered cytoplasmic puncta (Fig. S4B vs. Fig. 2d, 2e). These findings suggest that energy insufficiency can induce concentric cytoplasmic rings (CCR). These rings comprise stress granules, containing G3BP1 and nuclear RNA-binding proteins, that encircle the Golgi, which itself surrounds the aggresome together with VCP and Vimentin, and the centriole forms the nexus of the entire structure (Fig 2h).

To visualize the perinuclear concentric rings by an orthogonal approach, we applied transmission electron microscopy (TEM). In addition to the normal cellular content of control untreated cells, those treated with 2DG and Rotenone displayed membrane-less structures (Fig. S5A), consistent with the stress granules detected here by immunocytochemistry (ICC), and previously reported by others using the same technique (Bounedjah et al., 2014; Gilks et al., 2004). In the case of m.3243G, respiratory-deficient fibroblasts treated with 2DG, TEM also corroborated the ICC analysis, revealing stress granules in a ring, within which was the Golgi, and at its center a concentration of splayed cytoskeletal elements, which we infer is the aggresome (Fig. S5B), and which we never observed in any of our untreated control or patient-derived fibroblasts, in this or any previous study. On the other hand, no vacuoles or lysosomes were evident inside the ring of stress granules in the TEM images (Fig. S5B), nor were lysosomes redistributed when the CCR formed (Fig. S5B, S5C), suggesting that neither lysosomes nor vacuoles are CCR components.

### Repressed mitochondrial and cytosolic protein synthesis accompany the CCR response

Although energy insufficiency induces stress granules in primary human fibroblasts (Fig. 1, 2, S2, S3), m.3243G cells must possess some additional feature beyond respiratory chain deficiency to account for the induction of the cytoplasmic rings, as such rings were not seen in proliferating control fibroblasts treated with Rotenone and 2DG or KA. One possibility was impaired mitochondrial protein synthesis, as this process is disrupted by the defective transfer RNA (Leucine UUR) resulting from m.3243G (King et al., 1992). Therefore, we tested two inhibitors of mitochondrial translation, Chloramphenicol and Actinonin (the latter an inhibitor of peptide deformylase and several aminopeptidases), in combination with restricted glycolysis. 2DG, KA or glucose starvation with 150 µM Actinonin for 24 hours or KA with 50 µg/mL Chloramphenicol for 96 h, produced perinuclear rings of stress granules in a substantial majority (86%, 92%, 84%, 78%, respectively) of control fibroblasts (Fig. 3a, 3b, S6A-S6C), similar to those seen in m.3243G cells when glycolysis was restricted (Figs. 1b, S3). These findings suggest that impaired mitochondrial protein synthesis to the point of respiratory chain impairment triggers the CCR stress response in combination with impaired glycolysis. There is also a marked decrease in mitochondrial translation and respiratory chain capacity in (14 days) serum-deprived fibroblasts (Fig. 3c), and so we challenged such fibroblasts with 2DG and Rotenone and this induced the stress granules to form perinuclear rings, in approximately half of the cells (Fig. 3d, S6D); i.e., the arrangement seen in the control cells supplemented with serum and treated with Actinonin and 2DG (Fig. 3a). Although serum-deprivation and Actinonin arrest cell growth, other methods of inhibiting cell division did not induce stress granules or the CCR. First, 20 mM 2DG markedly inhibits cell growth and yet requires Rotenone or Actinonin to induce stress granules and the CCR, respectively, in control fibroblasts. Second, prior inhibition of the cell cycle with cyclin-dependent kinase inhibitor Dinaciclib did not affect stress granule distribution in 2DG/Rotenone treated cells (Fig. S6E-S6G). Thus, while energy scarcity will inevitably limit cell proliferation, and cessation of cell division is evident in conditions where the CCR forms, cell growth arrest is not the trigger of the CCR, and while inhibition of ATP producing pathways induces stress granules, it appears that the formation of the CCR additionally depends on inhibition of mitochondrial protein synthesis.

**Figure 3.**
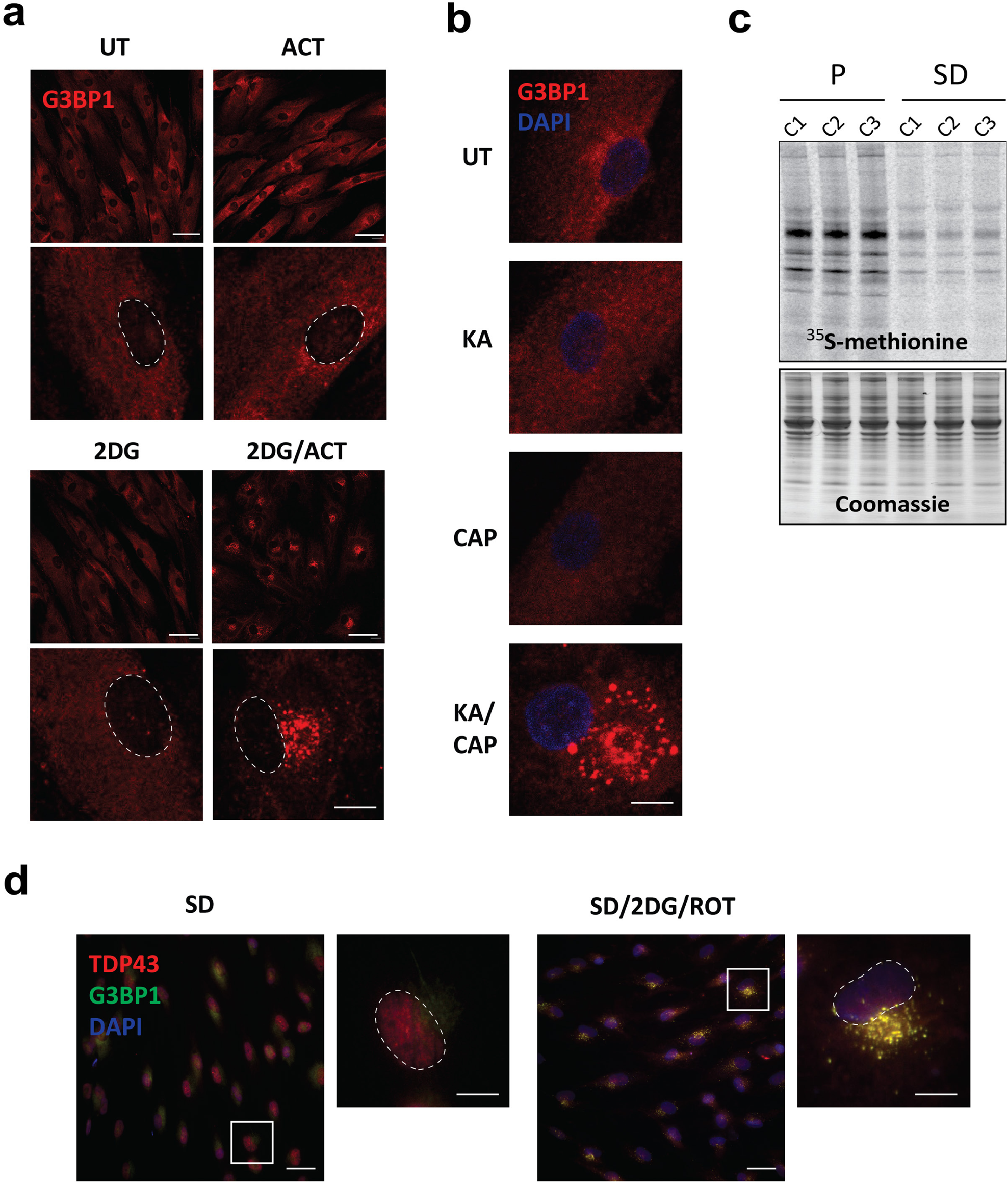
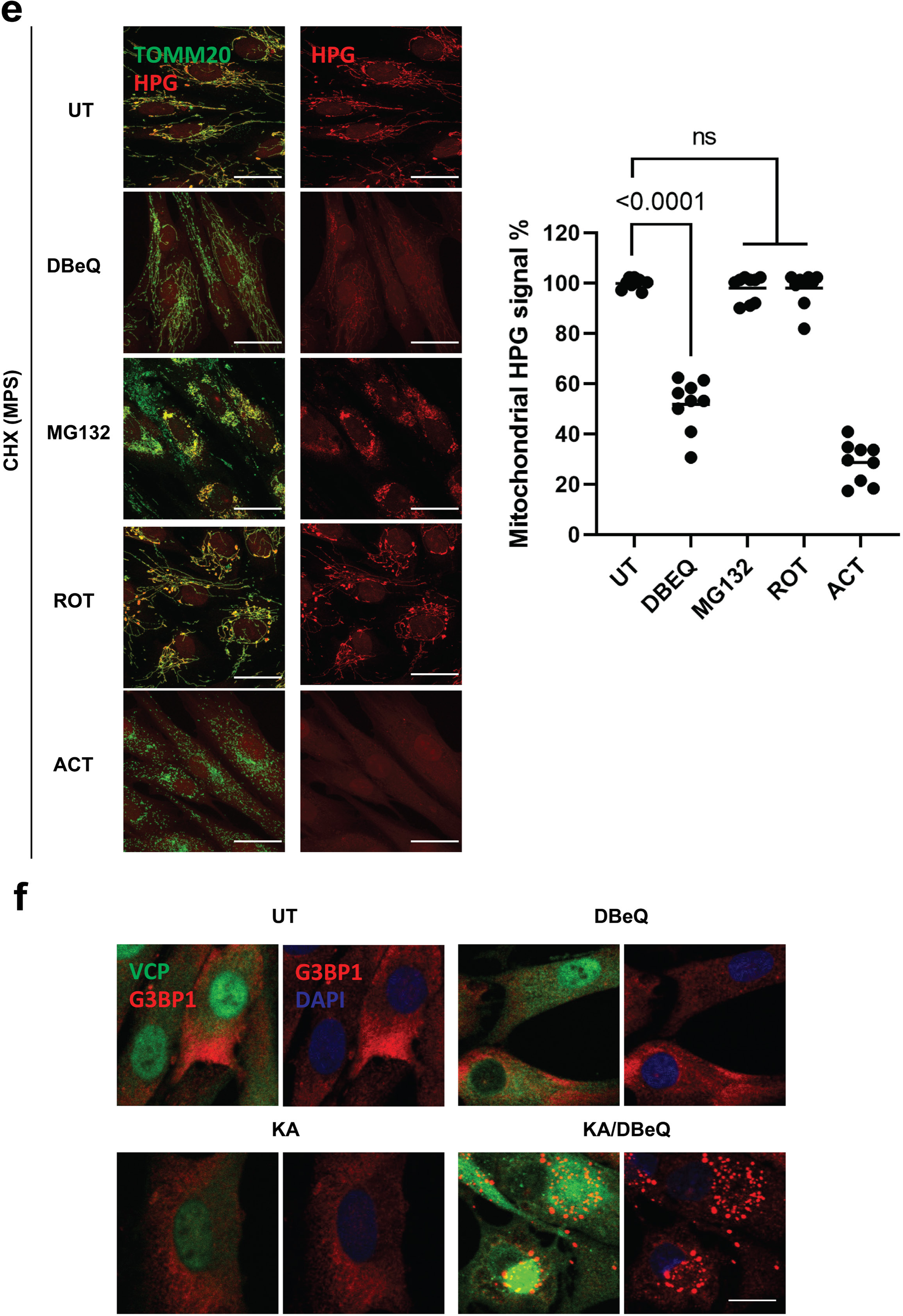
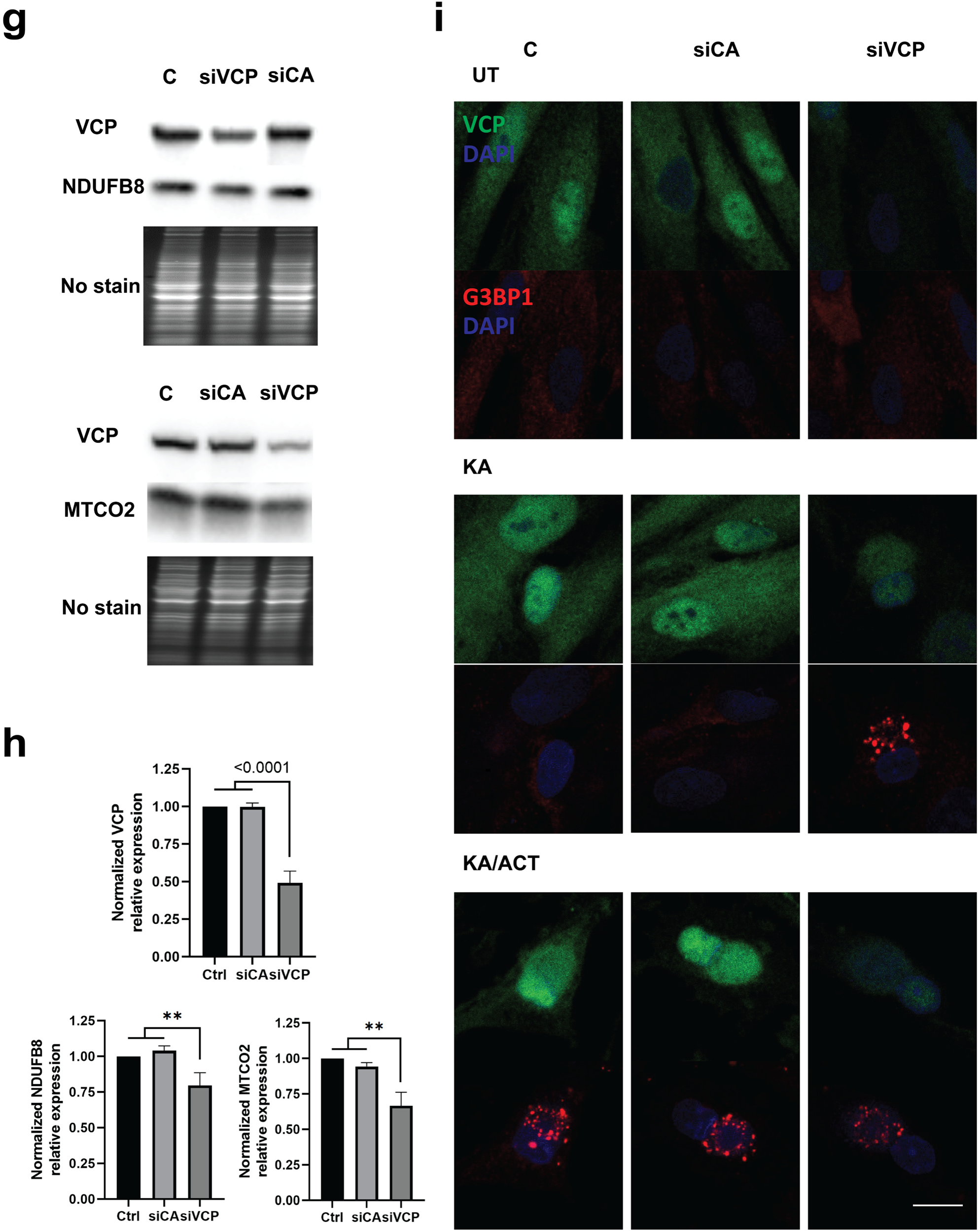
Impaired mitochondrial protein synthesis and serum restriction promote the induction of the CCR. **a**) Proliferating control fibroblasts were left untreated (UT) or treated with 150 µM Actinonin (ACT) and 20 mM 2DG for 24 h, as indicated, and immunostained for G3BP1 (red). Scale bar: 30 µm. Representative images of n=11 independent experiments in 5 control cell lines. **b**) As panel **a**, with treatments of 50 µg/mL Chloramphenicol (CAP), 1 µM Koningic acid (KA) and 20 mM 2DG, and additional staining of nuclei with DAPI (blue). Scale bar: 30 µm. Representative images of n=8 independent experiments on 3 control cell lines. **c**) Mitochondrial protein synthesis products were labeled with ^35^S-methionine in Emetine-treated proliferating (P) and serum-deprived non-dividing (SD) 3 control fibroblasts (C1, C2 and C3). Extracted protein was fractionated by SDS-PAGE and stained with coomassie blue (lower panel), to confirm equal protein loading, and exposed to a phosphor-screen to detect radiolabeled proteins (upper panel). **d**) After 14 days of serum deprivation (SD), control human fibroblasts were treated with 20 mM 2DG for 24 h and 0.1 μM Rotenone (ROT) for the final 3 h, and immunostained for G3BP1 (green) and TDP-43 (red). Nuclei are stained blue with DAPI. Scale bars: 30 µm. Representative images of n=10 independent experiments in 4 control cell lines. **e**) Fibroblasts, in which cytosolic translation was inhibited with 50 μg/mL Cycloheximide (CHX), were treated without (UT) or with 5 µM of the VCP inhibitor, DBeQ, or 5 µM of the proteasome inhibitor MG132, or 0.1 µM Rotenone, or 150 µM Actinonin, and in all cases the medium was supplemented with 50 μM of the methionine analog HPG to label nascent mitochondrial translation products. After fixing, HPG was detected by click chemistry with Alexa Fluor 555 azide (red) and the mitochondrial network was stained with TOMM20 (green). Scale bars: 50 µm. The chart to the right shows the HPG signal coincident with the mitochondria according to the different treatments. n=9 independent experiments in 3 control cell lines, a minimum of 270 cells analyzed in total for each condition. **f**) Fibroblasts were treated without (UT) or with 5 µM of the VCP inhibitor DBeQ, and additionally with (+) or without (-) 1 µM KA for 24 h, after which cells were immunostained for G3BP1 (red) and VCP (green). Scale bars: 30 µm. Representative images of n=9 independent experiments on 3 control cell lines. **g**) Immunoblot analysis of VCP, NDUFB8 (subunit of respiratory complex I), and MTCO2 (subunit of respiratory complex IV). Protein was extracted from human fibroblasts treated without (UT) or with silencer RNA targeting VCP (siVCP) or no target (siCA) for 72 h and exposed to 1 µM KA or 150 µM Actinonin for the final 24 h. A generic protein stain (No-stain^TM^) indicates relative protein loading. A representative blot from 4 independent experiments. **h**) Charts show the abundance of the 3 proteins in the different conditions from 4 independent experiments using 2 control cell lines. **i**) Fibroblasts immunostained for VCP and G3BP1, after no silencing (C) or siVCP or siCA without (UT) or with KA and Actinonin as indicated. Scale bar: 30 µm. Representative images of n=8 independent experiments on 2 control lines.

ALS-FTD is heavily associated with disturbed RNA metabolism, with consequences for protein production (Lehmkuhl and Zarnescu, 2018; Ling et al., 2013), and stress granule induction is associated with repressed cytosolic protein synthesis. Therefore, we assessed the impact of energy scarcity and mitochondrial dysfunction on cytosolic translation. Cytosolic protein synthesis is unaffected by Rotenone treatment but is repressed by 2DG, Actinonin or serum restriction, evidenced by reduced incorporation of an alkyne methionine analog, L-Homopropargylglycine (HPG), in cellular full-length proteins in three-hour pulse labeling experiments (Fig. S7A). Hence, partial inhibition of cytosolic, as well as mitochondrial, translation occurs in all the conditions that induce the perinuclear ring of stress granules. On the other hand, severe inhibition of cytosolic translation with Cycloheximide or Harringtonine abrogated stress granule formation in cells treated with 2DG and Rotenone or Actinonin (Fig. S7B, S7C).

VCP deficiency and mutation are associated with a range of mitochondrial abnormalities, especially mitochondrial protein turnover (Nalbandian et al., 2013; Tanaka et al., 2010; Xu et al., 2011), although there are contradictory reports as to its effect on the respiratory chain and mitochondrial membrane potential (Bartolome et al., 2013; Fang, 2015). Hence, we considered that altered VCP and mitochondrial function might be a crucial intersection in ring formation, owing to interdependence. And as perturbed mitochondrial translation appeared to be a core feature of CCR induction, we determined the effect of the selective and reversible inhibitor of VCP, DBeQ (*N^2^,N^4^*-dibenzylquinazoline-2,4-diamine) on translation. To specifically label mitochondrial translation products, cells were incubated with HPG in the presence of Cycloheximide (Yousefi et al., 2021), and the addition of Actinonin (or Chloramphenicol) reduced the signal to background, as expected, whereas Rotenone had no such effect (Fig 3e). Treatments of 5 µM DBeQ for 24 hours decreased mitochondrial protein synthesis 70% in a 3-hour pulse-labeling assay (Fig 3e), whereas parallel assays without Cycloheximide revealed a milder effect of DBeQ on cytosolic protein synthesis (a decrease of 21% compared to control levels) (Fig. S7D). As VCP mediates the unfolding and degradation of ubiquitylated proteins by the 26S proteasome, we tested the proteasome inhibitor MG132, in parallel with DBeQ. In striking contrast to DBeQ, 5 µM MG132 reduced cytosolic protein synthesis to a quarter of the control level, while having only a modest effect on mitochondrial translation (Fig. 3e and S7D). Thus, inhibition of VCP impedes mitochondrial translation independent of the proteasome. Next, we applied DBeQ in concert with KA, and the combined treatment induced the perinuclear ring of stress granules (Fig. 3f, S7E). Repressing VCP expression by RNA interference combined with KA also produced the perinuclear stress granule ring (Fig. 3g-3i, S7F), and reduced the level of respiratory chain proteins, MTCO2 and NDUFB8, by a third and a quarter, respectively (Fig. 3g, 3h). These findings indicate that VCP depletion or inhibition mimic Actinonin, Chloramphenicol and m.3243G, when glycolysis is inhibited (Figs. 1b and 3), and so suggest VCP perturbation of mitochondrial translation is a key reason it induces the CCR stress response. Equally, Actinonin, Chloramphenicol and m.3243G might inhibit VCP, when glycolysis is repressed, if mitochondrial translation and VCP activity are interdependent, and m.3243G might inhibit VCP, when glycolysis is repressed, if mitochondrial translation and VCP activity are interdependent.

### Selective Inhibition of nuclear protein export does not affect stress granule or CCR formation

Given that many stress granule components ordinarily function as nuclear RNA binding proteins, a change in nuclear protein transport owing to energy scarcity might contribute to (perinuclear) stress granule formation. Therefore, we tested a selective inhibitor of nuclear protein export (SINE) - Verdinexor (KPT-335), a specific XPO1/CRM1 inhibitor. KPT-335 treatment (150 nM, 24 h) caused SQSTM1 to accumulate in the nucleus (an 85% increase in nuclear signal) and increased the nuclear FUS signal 55% (Fig. S8A, S8B), suggesting that nuclear protein export was blocked as intended, whereas nuclear protein import was maintained. However, an overnight pre-treatment with KPT-335 did not prevent relocalization of FUS to the cytoplasm when cells were treated with 2DG and CCCP for 3 hours, as there was no significant difference in residual nuclear FUS signal with and without the KPT-335 (Fig. S8A). Nor did KPT-335 diminish stress granule formation, or prevent, or induce CCR formation (Fig. S8B). These results suggest that nuclear protein import and export are independent of stress granule and CCR formation, and so we tentatively infer that the nuclear RNA binding proteins of the stress granules exit the nucleus by passive diffusion when energy is scarce, as suggested by others (Duan et al., 2022; Ederle et al., 2018).

### Induction of the CCR is associated with inhibition of autophagy whereas ERAD and proteasome activity are maintained

We previously found that autophagy is inhibited when 2DG is applied to respiratory deficient cells (with high levels of m.3243G), or to control cells in combination with Rotenone (Pantic et al., 2021), i.e., the autophagy system is inhibited when dispersed or perinuclear stress granules are induced by energy restriction. However, inhibition of autophagy is not sufficient for stress granule or CCR formation, as neither was induced in control fibroblasts by Chloroquine treatment, alone or in combination with 2DG, or Rotenone (Fig. S9A). Moreover, while Chloroquine with 2DG induced VCP-containing aggregations, these were large, amorphous and dispersed, unlike the unique CCR foci (Fig. S9A vs. Fig. 2).

The core of the CCR, the aggresome, forms in some cells in response to proteasome inhibition, and can (Garcia-Mata et al., 1999; Johnston et al., 1998; Shin, 1998), but need not (Matsumoto et al., 2018), contain misfolded ubiquitinated proteins. However, while treatments of human fibroblasts for 24 hours with 5 µM of the proteasome inhibitor MG132 induced large aggregations of VCP and SQSTM1 in control and respiratory deficient fibroblasts (Fig. 4a), these were quite different from the aggresome and the CCR. First, the MG132-induced inclusions were numerous and widely dispersed in most of the cells; second, they contained the full cytoplasmic complement of SQSTM1, which was non-phosphorylated; third, there was no Vimentin cage, nor fourth, cytoplasmic granules of G3BP1/TDP-43 (Figs. 4a, S9B). Therefore, inhibition of the proteasome produces inclusion bodies, but does not induce the aggresome or stress granule formation in primary human fibroblasts. The clear inference that CCR formation is not attributable to proteasome inhibition was corroborated by tests that showed energy scarcity reduced proteasome activity little in proliferating cells, and still less in serum-deprived cells (Fig. S9C). Although many misfolded proteins are fed to the proteasome via the Endoplasmic-reticulum-associated protein degradation system (ERAD), often in conjunction the unfolded protein response (UPR), neither correlated with the appearance of the stress granules or the CCR. While the UPR was elevated in proliferating control cells treated solely with 2DG, it was considerably lower in cells treated with 2DG and Rotenone, and equally low in m3243G or serum-deprived cells treated with 2DG, and not evident at all in cells treated with KA (Figs. 4b, 4c, S9D, S9E, S9G). To evaluate ERAD, we assessed CD147 an endogenous substrate of the system that is degraded via the Hrd1/SEL1L pathway (Tyler et al., 2012). CD147 has a plasma membrane form with complex glycans (CD147 Mat), and an ER form bearing the core-glycan structure (CD147 CG) (Tang et al., 2004). While CD147 processing was impaired by inhibition of N-glycosylation with 2DG, Tunicamycin, or the ERAD inhibitor Kifunensine (Elbein et al., 1990), it was not affected by KA, with or without Actinonin, Rotenone or Chloroquine (Fig. S9E-G). Hence, neither stress granules nor the CCR are induced in response to UPR induction or ERAD dysfunction.

**Figure 4.**
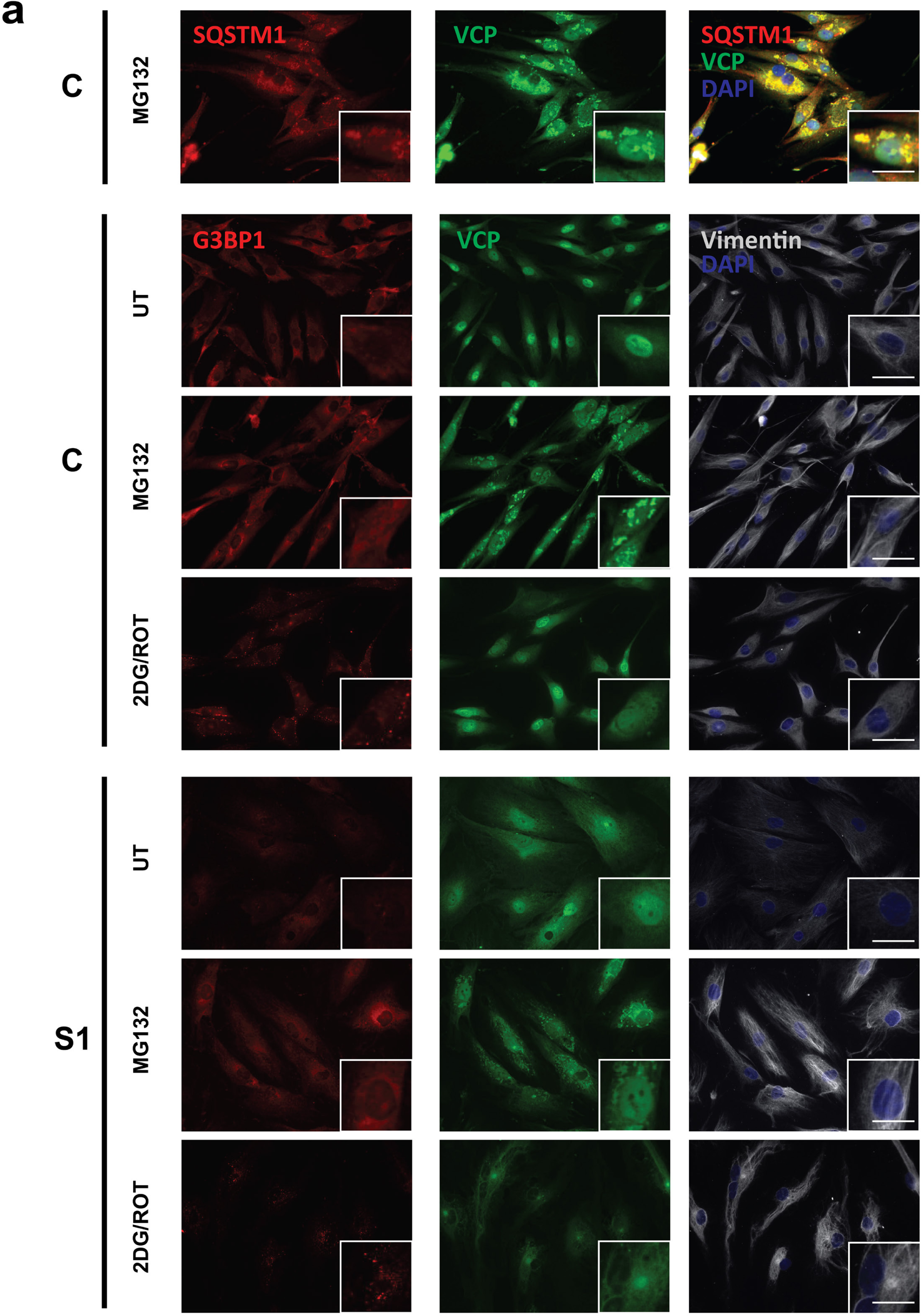
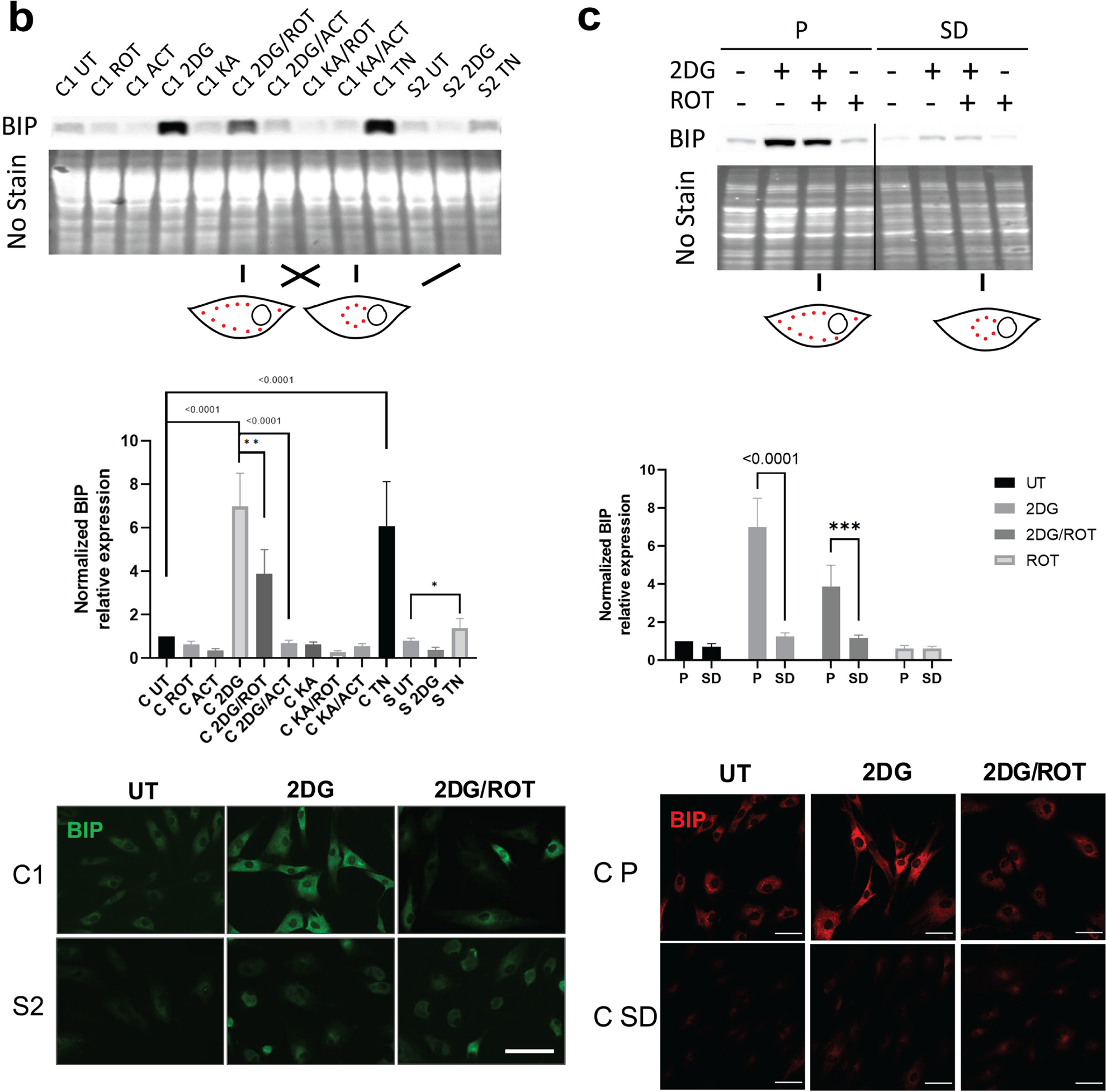
The CCR response is independent of proteasome inhibition, and UPR induction, but is associated with inhibition of autophagy. **a**) Control fibroblasts treated with 5 µM MG132 for 24 h and immunolabeled for VCP and SQSTM1 (upper row). Control and respiratory deficient (S1, m.3243G) fibroblasts treated with 5 µM MG132 for 24 h, or 24 h with 20 mM 2DG and for the final 3 h with 0.1 µM Rotenone, and fixed and immunolabeled for G3BP1, VCP, and Vimentin (lower rows). Scale bar: 50 µm. Representative images of 4 independent experiments using 2 mutant cell lines and 4 control cell lines. BIP (GRP78) protein expression assessed by immunostaining in **b**) control (C) and respiratory deficient cells (S2, m.3243G) and **c**) control proliferating and serum deprived fibroblasts; treatments 2DG, 1 µM KA, 1 µM Tunicamycin, (TN) and 0.1 µM rot (Rotenone); total protein was detected by No stain^TM^. Scale bar: b) 100 μm and c) 50 μm. b) and c) Blots and images are representative from 4 independent experiments with 4 control cell lines and b) S2 mutant cell line. Illustrations below the blots indicate the occurrence of stress granules (red dots) towards the cell periphery, or the perinuclear ring of stress granules.

### Microtubule depolymerization impairs the formation of the CCR, whereas microtubule acetylation induces stress granules to form a perinuclear ring

Several factors pointed to microtubules being a potential key factor for CCR formation and maintenance. The microtubule organizing center lies at the center of the CCR, and the aggresome surrounding it is a microtubule-dependent entity (Kopito, 2000), to which stress granules can be trafficked for recycling in some circumstances (Kawaguchi et al., 2003; Kwon et al., 2007; Mateju et al., 2017). Therefore, we applied an inhibitor of microtubule polymerization, Nocodazole, to determine the contribution of the microtubule network to the CCR. While stress granules (labeled with G3BP1 and FUS) formed perinuclear rings as previously when control fibroblasts were treated overnight with KA or 2DG and Actinonin, co-treatment with 5 µM Nocodazole reduced the normalized diameter of the aggresome approximately four-fold, from 0.28 to 0.08, and dispersed the G3BP1 signal, reducing its perinuclear concentration more than five-fold, from 68% to 12% (Figs. 5a and S10A).

**Figure 5.**
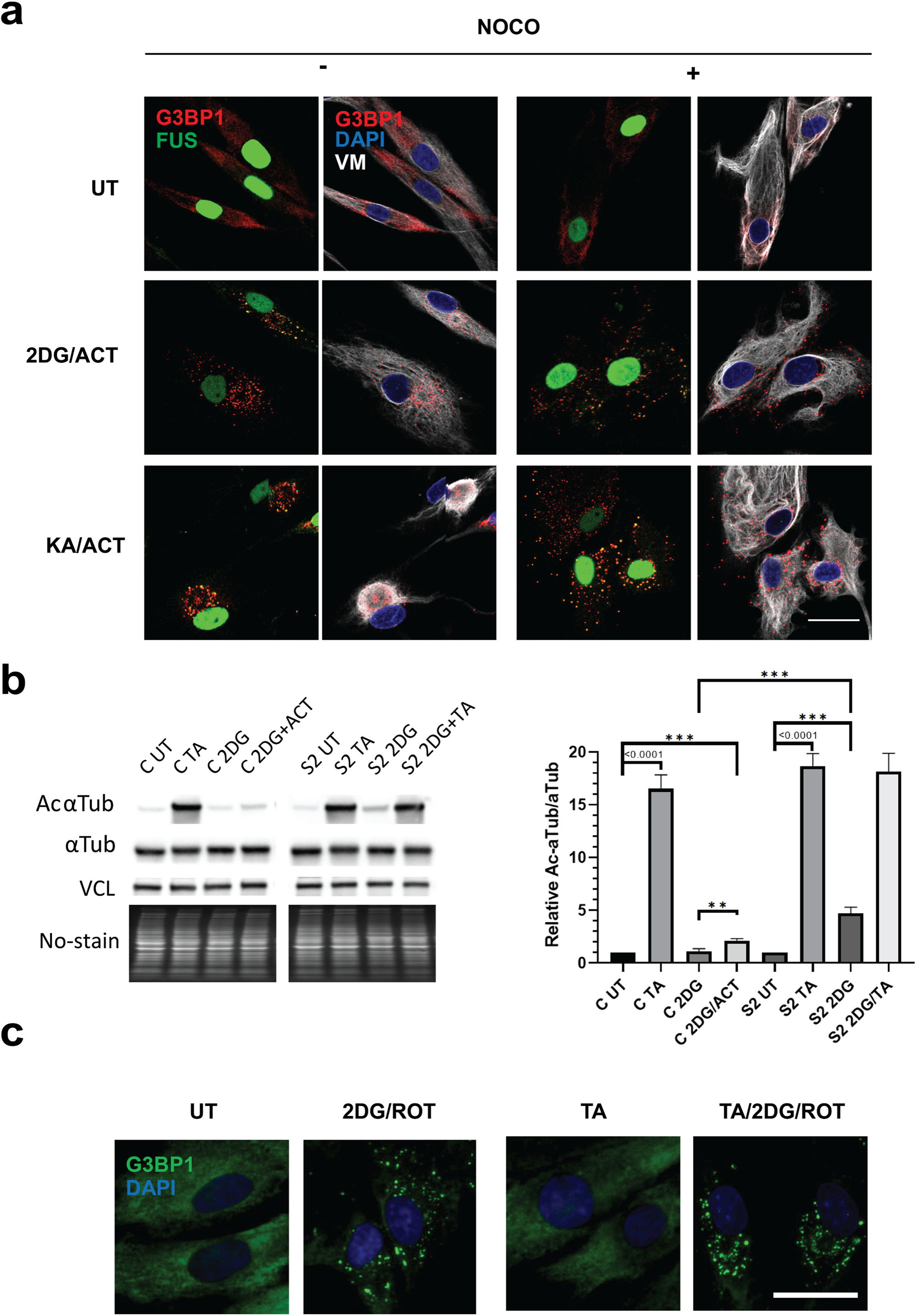
Disruption of the microtubule network disrupts the perinuclear ring of stress granules whereas HDAC6 inhibition causes dispersed stress granules to form a perinuclear ring. **a**) Control fibroblasts treated without (UT) or with 20 mM 2DG or 1 µM KA and 150 µM Actinonin for 20 h, all with (+) or without (-) 5 µM Nocodazole, and fixed and immunolabeled for G3BP1, and FUS. Scale bar: 100 µm. Representative images from the analysis of 270 cells, across n=9 independent experiments on 3 control fibroblast lines. **b**) Immobilized proteins immunostained for α-tubulin, acetylated α-tubulin and the loading control vinculin and No-stain as another loading control from control human fibroblasts or respiratory deficient cells (S2, m.3243G). Cells were untreated (UT) or exposed for 24 h to one or more of the following: 20 mM 2DG, 150 µM Actinonin (ACT), 10 µM Tubastatin A (TA). On the right, the chart displays the quantification of n=3 independent experiments using 3 control fibroblasts and the S2 mutant cell line. **c**) Control fibroblasts were left untreated (UT), or exposed to 0.1 μM Rotenone for the final 3 h of a 24 h, 20 mM 2DG treatment, with (+) or without (-) TA. After fixing, cells were immunostained for G3BP1. Representative images from n=9 independent experiments on 3 control cell lines and analyzing a total of 450 cells per condition. Scale bar: 50 µm.

Stress granule formation is mediated by the motor-protein-driven movement of individual components along microtubules, and this process is modulated or mediated by the deacetylase HDAC6 (Kwon et al., 2007), and the same transport system recruits and delivers misfolded proteins to the aggresome (Kawaguchi et al., 2003). Therefore, we analyzed the effect of the HDAC6 inhibitor Tubastatin A (TA) on the dispersed stress granules, after confirming that 24 hours exposure to 10 µM TA increases α-tubulin acetylation 16-fold in fibroblasts (Fig. 5b). The inhibition of HDAC6 with TA in fibroblasts treated with a combination of 2DG and Rotenone caused the stress granules to form a single perinuclear ring in 89% of cells (Fig. 5c, S10B). This finding suggests that HDAC6 traffics stress granules (or deacetylates α-tubulin to permit stress granule trafficking) towards the cell periphery. As a corollary, we infer that HDAC6, or an ancillary factor, is inhibited under the conditions where the CCR forms. Furthermore, there was a 100% increase in acetylated α-tubulin when control cells were treated with 2DG and Actinonin, and a five-fold increase when m.3243G fibroblasts were treated with 2DG alone, indicating that microtubule acetylation is elevated in the contexts where the CCR forms (Fig. 5b).

### Perinuclear stress granules are soluble, but more stable than dispersed stress granules, and their dissolution is dependent on the restoration of glycolysis

The inclusions of ALS-FTD, and other neurodegenerative diseases, are essentially insoluble, irreversible structures, whereas stress granules dissolve within minutes or hours (Hofmann et al., 2021). Hence, stress granules must both coalesce and become insoluble if they are to be the progenitors of the perinuclear inclusion found in ALS-FTD. Thus, the concept of the aberrant stress granule has been forwarded as a potential key precursor of the insoluble inclusion (e.g. (Mateju et al., 2017); however, it is not clear how such aberrant structures form in sporadic forms of ALS. The dispersed stress granules and those of the CCR share a similar electron density in the TEM images (Fig. S5A, S5B), and they are soluble in 1,6-Hexadienol (1,6-HD), which dissolves liquid–liquid phase separated condensates (Kroschwald et al., 2017; Lindström et al., 2022) (Fig. S11A). Cessation of the 2DG and Rotenone treatment resulted in the dissolution of the peripheral stress granules within 20 minutes, whereas the perinuclear stress granules took 3 hours to disperse (Fig. 6a-6c, S11B, S11C). Hence, the stress granules of the CCR are almost an order of magnitude more stable than those more widely dispersed stress granules (Fig. 6c), a similar difference has been seen for cold-shock induced stress granules (Hofmann et al., 2012) versus other treatments such as heat-shock and Sodium arsenite (Hofmann et al., 2021). Furthermore, the (ring of) stress granules persisted when KA and Actinonin or KA and RRotenone were removed from the growth medium (Fig. 6b, 6c), suggesting that once triggered the stress granules and the CCR, they persist as long as glycolysis is repressed, as KA irreversibly inhibits glyceraldehyde 3-phosphate dehydrogenase (Endo et al., 1985).

**Figure 6.**
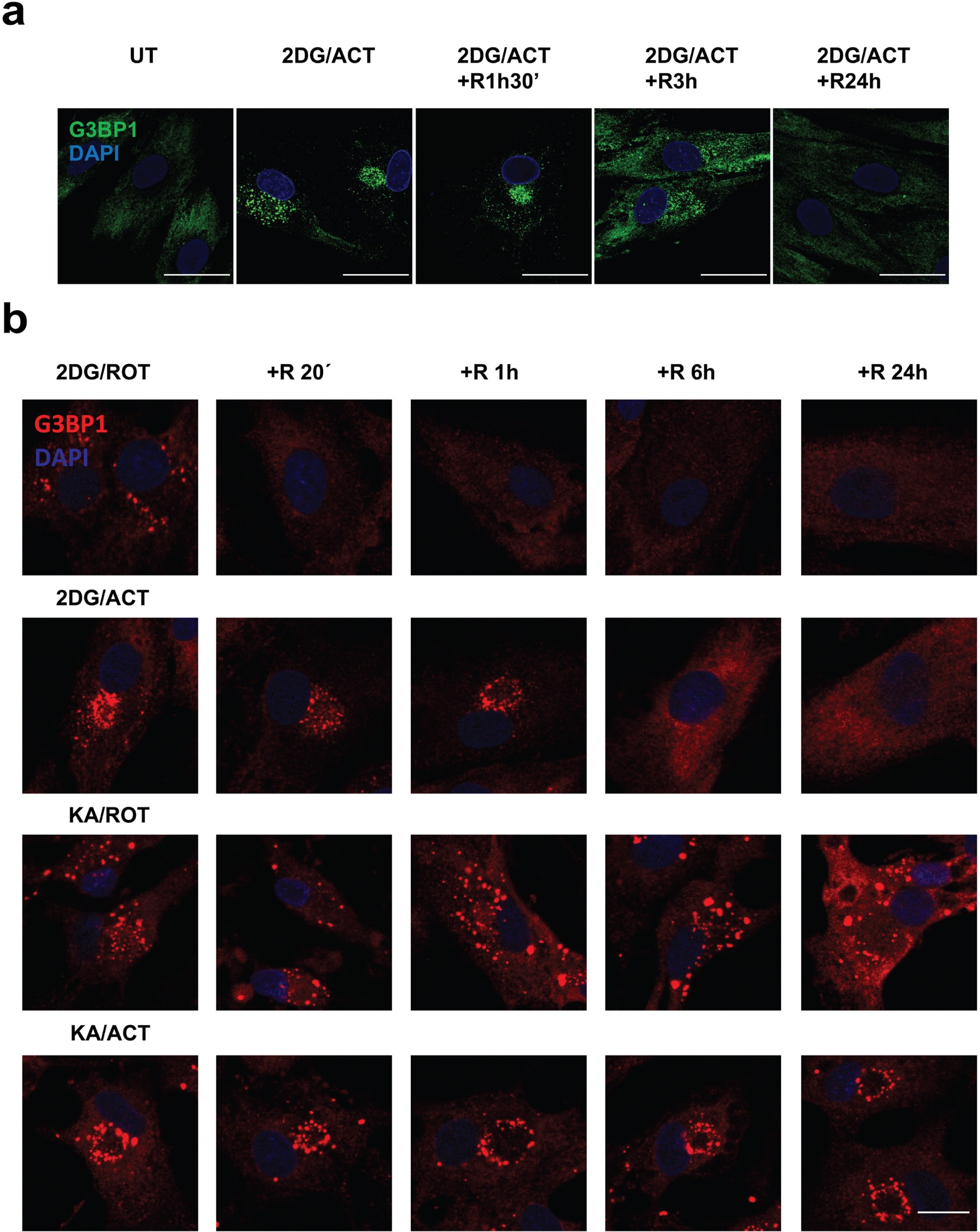
The perinuclear rings of stress granules are reversible. **a**) Human fibroblasts were untreated (UT) or subjected to 2DG and Actinonin (ACT) for 24 h, after which the compounds were removed and the cells stained with antibody to G3BP1 (green) and DAPI (blue), after no recovery or recovery periods (R) of 1.5, 3 and 24 h. **b**) Cells were treated for 24 h with 2DG and Rotenone (ROT), 2DG/ACT, Koningic Acid (KA)/ROT or KA/ACT and fixed and stained for G3BP1 (red) and DAPI (blue) after no recovery or recovery periods (R) of 20 minutes, 1, 6 and 24 h. Scale bars: 30 µm. Representative figures of n=6 independent experiments on 2 control cell lines. **c**) Chart indicating the time taken for stress granules to disappear (recovery time) after removal of the different inhibitors of the energy producing pathways: 2DG/ROT (orange), 2DG/ACT (pink), KA/ROT (green), KA/ACT (turquoise) (percent of cells with stress granules (SGs) or rings measured in 500 cells across n=10 independent experiments on 3 control cell lines).

To visualize the formation of the CCR in real time, we transfected cells with CellLight™ Golgi-GFP (Invitrogen) to express Golgi resident protein (N-acetylgalactosaminyltransferase) tagged with GFP and treated them with 2DG and Actinonin. Live cell imaging revealed a remodeling of the organelle into a ring over the course of 2-3 hours (Fig. S12A), consistent with immunolabeling of the Golgi in other experiments (e.g., Fig. 2a, S4A). On the other hand, a G3BP1-GFP fusion protein induced spontaneous stress granule formation, in approximately 50% of transfected cells, and extensive cell death without any accompanying treatment (Fig. S12B). Under conditions that induce the CCR, the surviving cells expressing the recombinant protein lacked the stress granules ring, whereas the perinuclear stress granules ring formed as expected in adjacent untransfected cells with no GFP signal (Fig. S12B). We also attempted to label stress granules with a small-molecule probe, TASG (Shao et al., 2021) without success (Fig. S12C).

### 2DG induces cerebral perinuclear concentrations of TDP-43/FUS reminiscent of the pathological hallmark of ALS-FTD

The brain is the major consumer of glucose; and so restricted glycolysis would be expected to be more challenging in the brain than in cultured fibroblasts; moreover, murine cells appear to be more sensitive to energy restriction than human fibroblasts ((Wang et al., 2019b) vs. this report). To determine whether 2DG was sufficient to induce protein translocation *in vivo* and create stress granules and structures akin to the CCR, we administered 0.4% w/v 2DG in drinking water three consecutive days a week for 8 months, to the standard laboratory mouse strain, C57BL/6J, which equated to approximately 1200 mg/Kg/week. Brain sections of six 2DG treated and six untreated controls were analyzed via immunolabeling. Immunostaining for phosphorylated (Ser409/410) TDP-43 (pTDP-43) revealed occasional small puncta in some cells of control mice (Fig. 7a), including when the primary antibody was omitted (Fig. 7b), but uniquely in the 2DG treated group perinuclear rings of pTDP-43 foci were detected, some more compact than others (Fig. 7c vs. 7d). Moreover, these rings coincided with cytoplasmic FUS, whereas in all the cells lacking such a ring, FUS was exclusively nuclear localized (Fig. 7a-d). Staining for specific cell types indicated that the pTDP-43 positive foci were associated with activated microglia (IBA1), where they accounted for half of the population (Fig. 7e), whereas none were detected in neurons (NeuN), astrocytes (GFAP), or oligodendrocytes (SOX10). The diameter of pTDP-43/FUS perinuclear ring was 3.3 ±0.3 µm in the activated microglia; while this was considerably smaller than that of the human fibroblasts, the diameter normalized to the nucleus was 0.55, similar to the normalized Dr of 0.64 in the fibroblasts (Fig. 7f). Hence, we infer that the 2DG-induced cellular structures in the microglia are stress granules arranged as per the CCR of the human fibroblasts (Fig. 7g). The health of the mice at the end of the 8-month intermittent 2DG treatment was not investigated in detail, but there were no deaths among the treated mice and no overt signs of motor or other neurological dysfunction.

**Figure 7.**
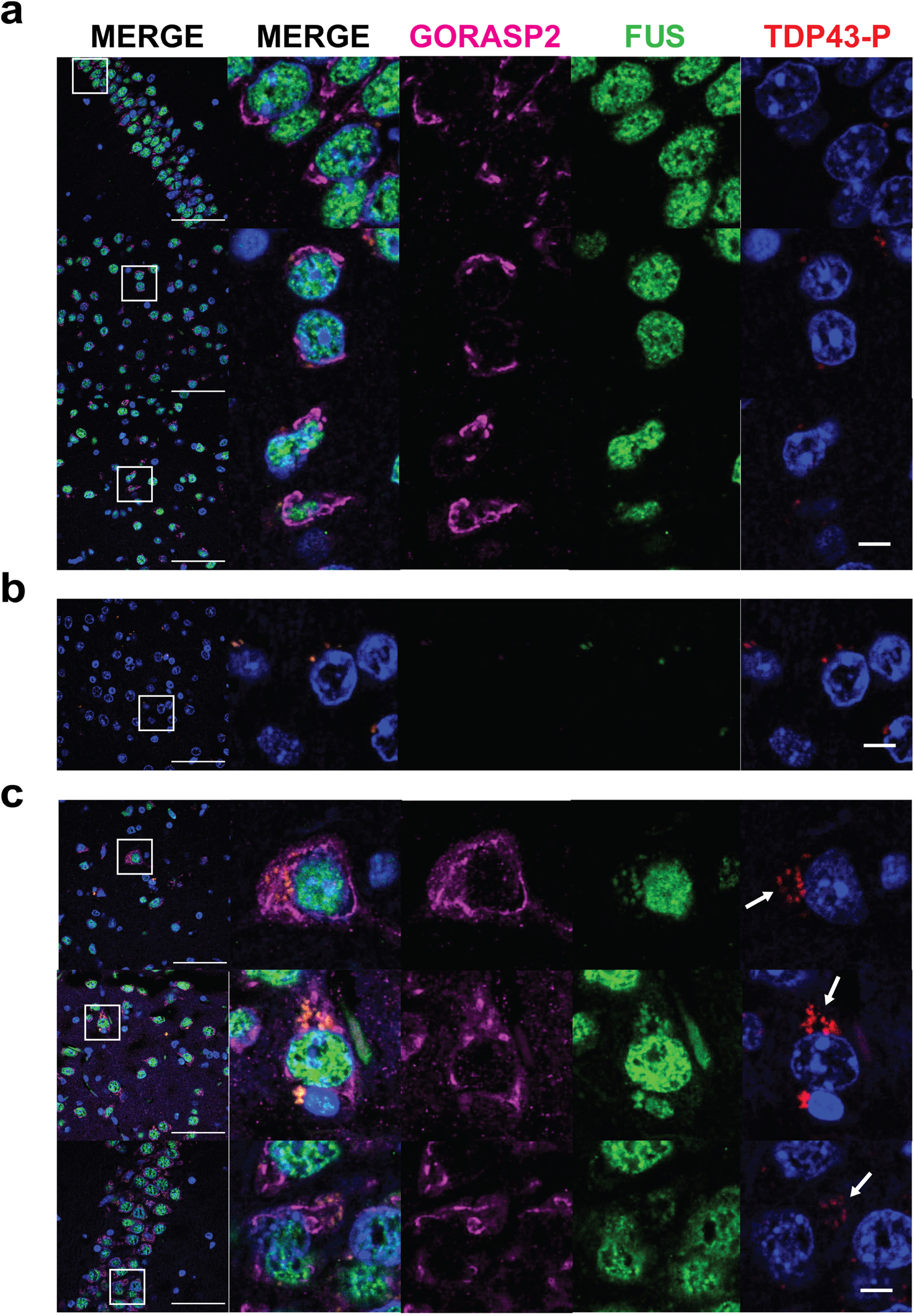
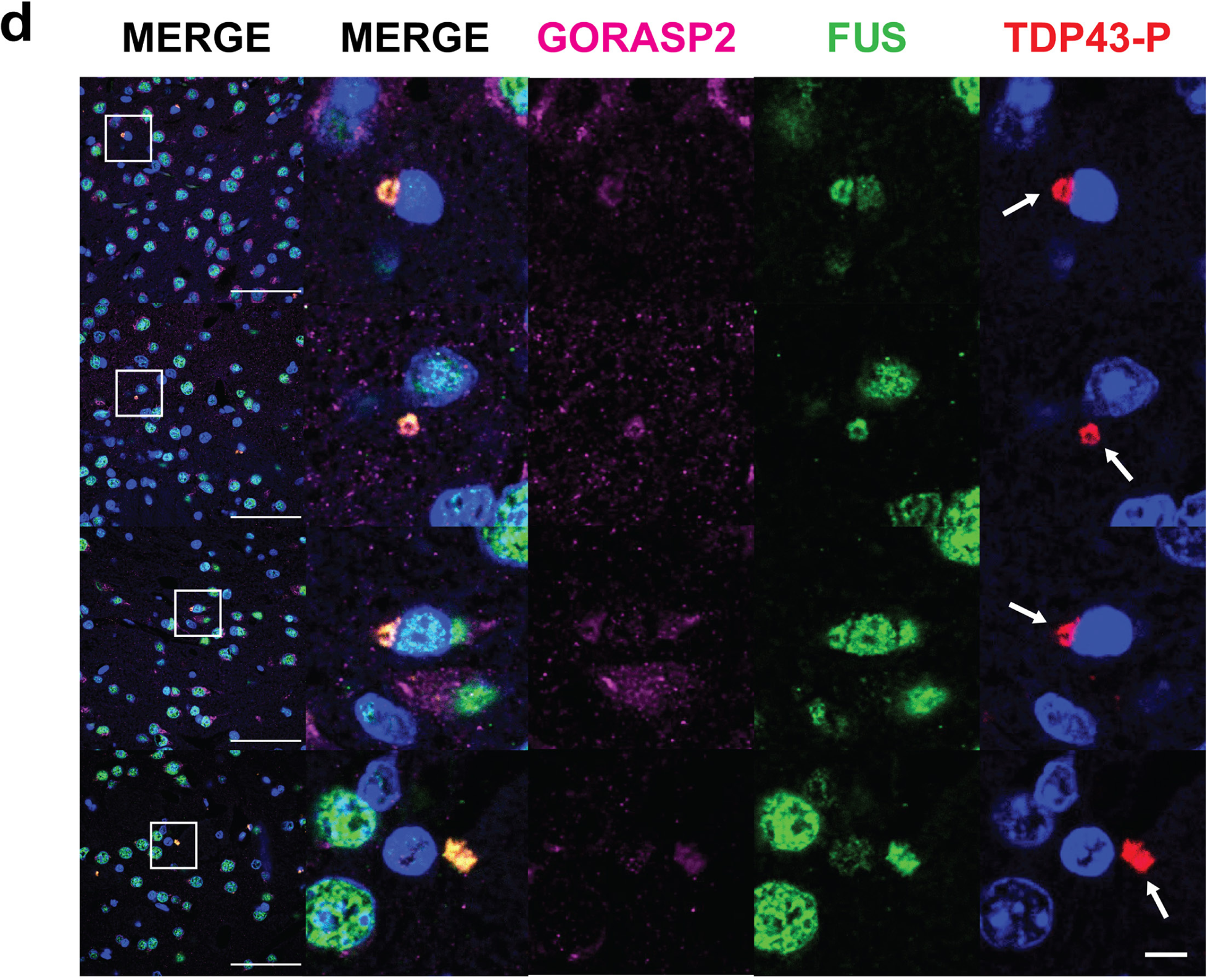
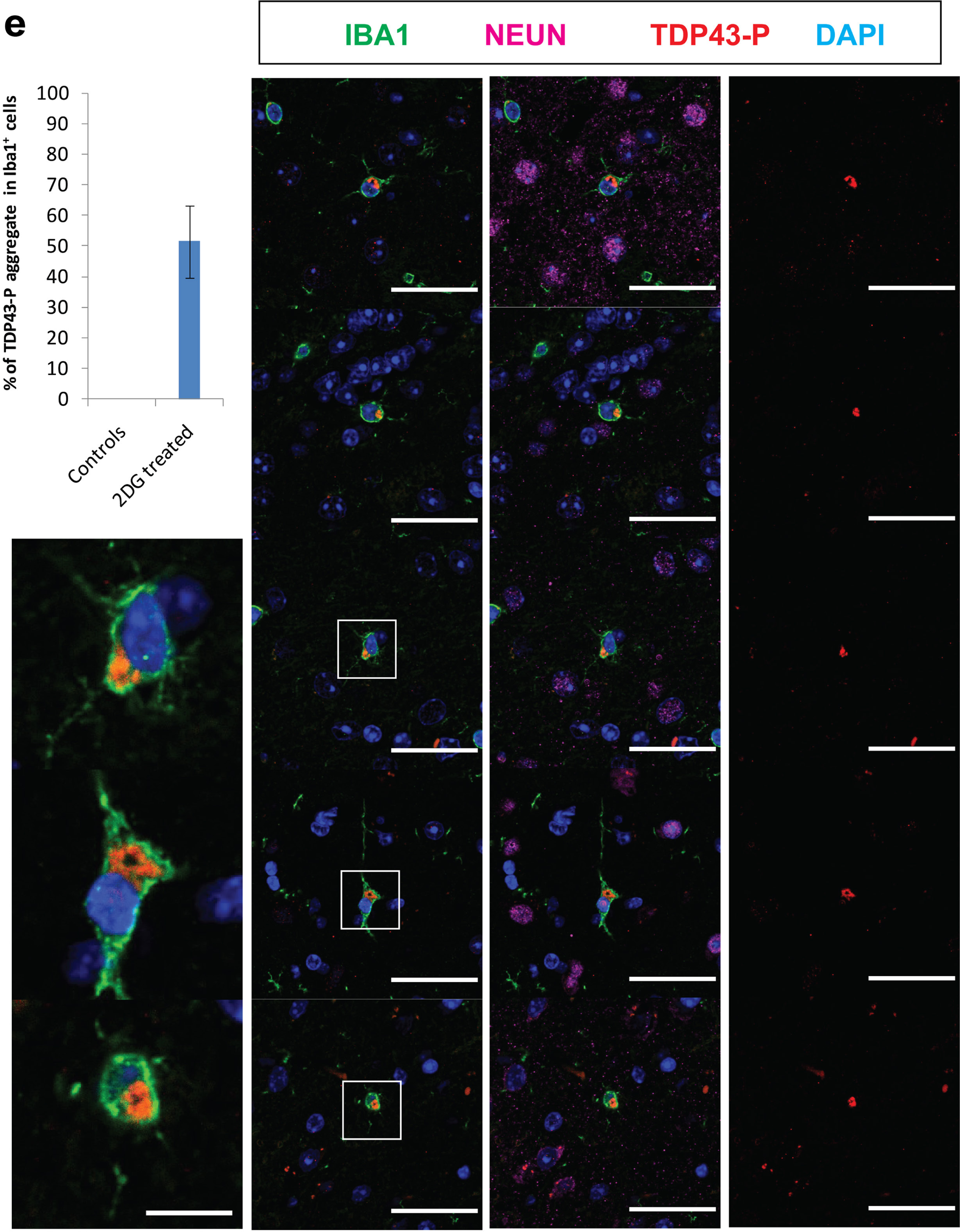
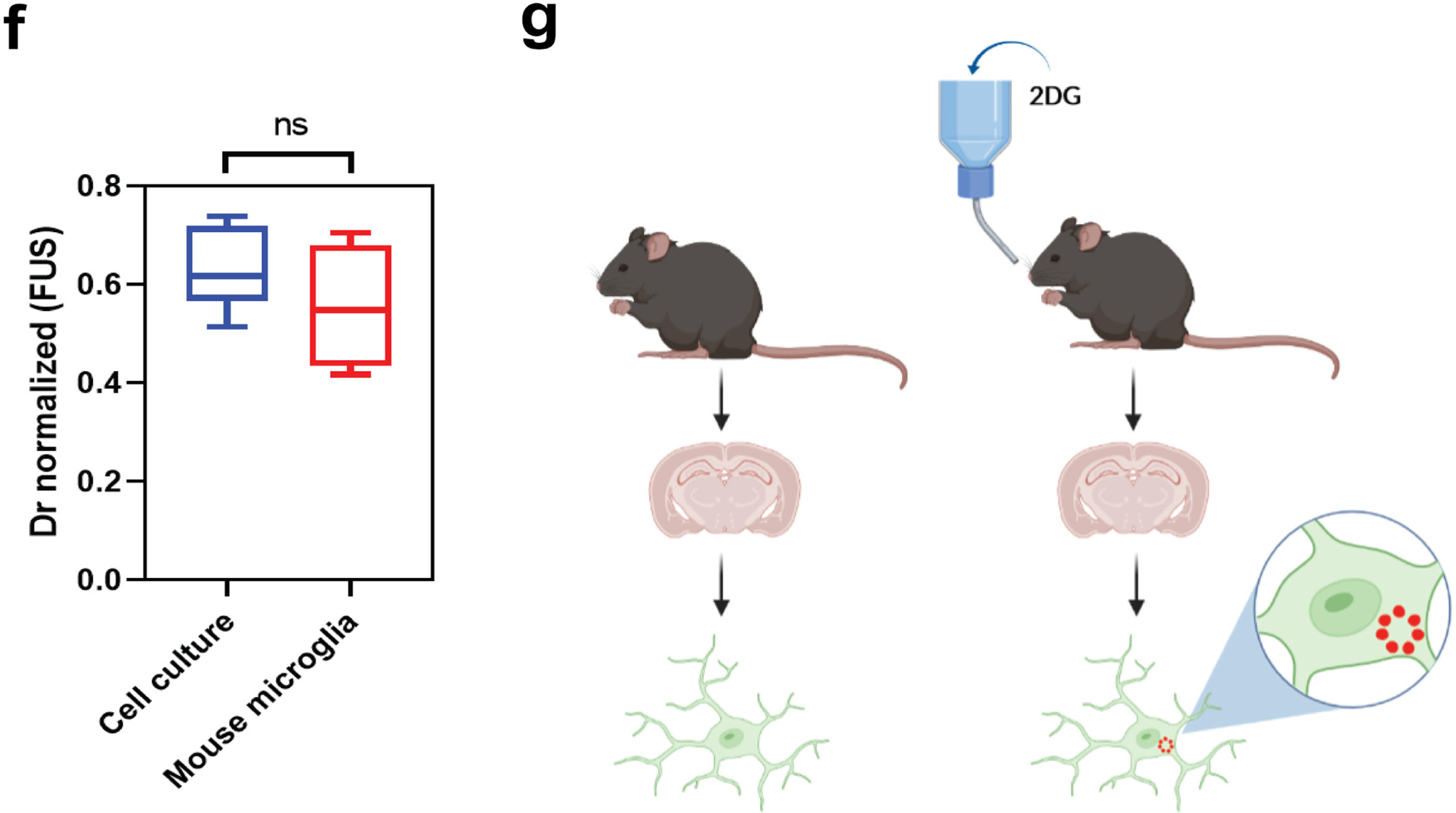
Perinuclear rings of stress granules form in the brains of mice treated with 2DG. Mice were treated without or with 0.4% (w/v) 2DG in the drinking water for three consecutive days per week for 8 months. Six treated and six untreated mice brain sections were immunostained in 3 independent experiments for each antibody. **a**) Fixed brain sections of untreated mice immunostained for phosphorylated (Ser409/410) TDP-43 (TDP43-P), FUS, GORASP2, the neuron marker NeuN, and the microglial marker IBA1; nuclei are stained blue with DAPI. To the left, fields of cells from sections of untreated murine brain (scale bar: 50 µm), and to the right higher magnifications (scale bar: 5 µm). **b**) Brain section of a treated mouse immunostained with the secondary antibody without a primary antibody, with DAPI stained nuclei (blue). **c**) and **d**) as panel **a**, fields of cells from sections of murine brain from animals treated with 2DG (scale bar: 50 µm), and to the right higher magnifications (scale bar: 5 µm) showing cytoplasmic rings of TDP43-P coinciding with FUS (i.e., stress granules). White arrows indicate the position of the ring. **e**) Murine brain sections treated with 2DG were stained for TDP43-P (red) and NeuN (purple), IBA1 (green) and DAPI (blue). To the left, zooms showing individual Iba1-expressing cells with a TDP43-P ring, and above the zooms a chart indicating the proportion of Iba1 stained cells with a TDP43-P ring from brain sections of 6 control and 6 2DG-treated mice (50 IBA1 positive cells were analyzed). Scale bar: 50 µm and in the zooms 12.5 µm. **f**) chart indicating the relative diameter of the stress granule rings in murine microglia and human fibroblasts, both normalized to the nucleus based on an analysis of a minimum of 150 microglia and 150 fibroblasts. **g**) Schematic representation of the 2DG treatment of mice that induces a perinuclear ring of TDP43-P (red dots) in active microglia (green cells).

## DISCUSSION

Many studies of ALS-FTD focus on models that carry mutant variants of proteins that relocate to the cytoplasm and form inclusions in the disease state, such as FUS and TDP-43. The limitation of these studies is that they do not explain how non-mutant forms of these proteins translocate and form a discrete perinuclear inclusion, as occurs in most sporadic cases. In this study, we show that dual inhibition of ATP producing pathways, encompassing impaired mitochondrial translation, induces a suite of ALS-associated factors to coalesce in concentric cytoplasmic rings in cultured cells. These rings include stress granules containing FUS and TDP-43 that are frequently found in a perinuclear body in ALS-FTD (Mori et al., 2008; Shintaku et al., 2017). Thus, perturbed nutrient and energy metabolism could generate an intermediate stage in the development of sporadic ALS-FTD. Moreover, the fact that microtubule acetylation via inhibition of HDAC6 causes stress granules to concentrate at the aggresome, rather than close to the cell periphery (Fig. 5), suggests a model for the development of the perinuclear body. Energy scarcity induces stress granules, and coupled with impaired mitochondrial translation, remodels microtubules causing stress granules to cluster around the aggresome (Figs. 2, 5 and S10). Furthermore, while the rings are triggered when both glycolysis and mitochondrial function are compromised, the stress response is maintained while glycolysis is repressed (Fig. 6c), which suggests transient mitochondrial impairment combined with chronic glycolytic insufficiency could create irreversible stress granules that evolve to inclusions and thus ALS-FTD (Fig. 8). The perturbation of mitochondrial function could evidently take the form of VCP inactivation, as VCP silencing or chemical inhibition, combined with repressed glycolysis induces the perinuclear ring of stress granules (Fig. 3).

**Figure 8.**
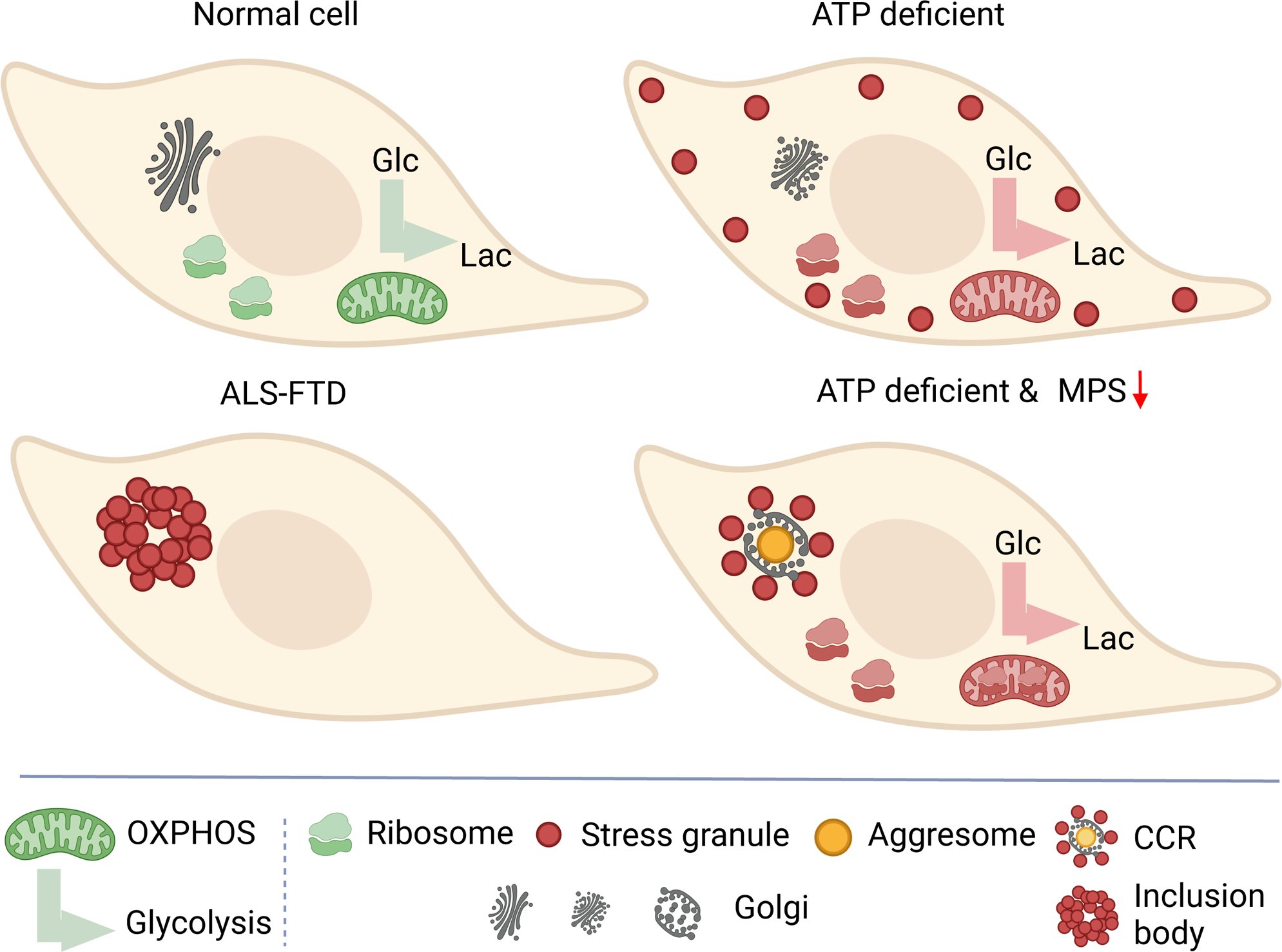
Model of the development of ALS-FTD associated inclusions owing to energy scarcity. In a normal healthy cell (top left), ATP is generated via the conversion of glucose to lactate (glycolysis) and in the mitochondria via the respiratory electron transport chain (RETC). Inhibiting these ATP producing pathways in cells induces relocation of a variety of nuclear proteins, such as TDP-43 and FUS, to the cytoplasm where they form stress granules that are spaced far apart towards the cell periphery (top right). In cells with low or without mitochondrial protein synthesis (MPS) the same treatment causes the stress granules to cluster in a perinuclear ring, surrounding a remodeled Golgi apparatus that encompasses the aggresome, aka the CCR (lower right). The CCR is formed in proliferating control cells subjected to glucose restriction and MPS inhibitors, and in glucose-restricted cells with a genetic defect in MPS owing to mutant mtDNA (m.3243G). Hence, stress granules form in response to partial ATP depletion and are more closely packed in conditions of reduced mitochondrial protein synthesis in primary human fibroblasts. We propose that further concentration of the stress granules leads to liquid to solid phase transition resulting in TDP-43 positive, insoluble inclusions that are the pathological hallmark of ALS-FTD; and that something similar occurs in activated microglia *in vivo*, when mice are treated with 2DG (bottom left).

Stress granule induction and ALS-FTD are heavily associated with disturbed RNA metabolism and cytosolic translation (Ling et al., 2013), and our findings highlight the impact of the mitochondrion on these processes. Actinonin inhibits translation in both compartments, which may stem from aberrant N-terminal peptides that in bacteria are associated with inclusion body formation (Király et al., 2006). Hence, it will be of interest to learn if this process is enhanced or accelerated by nutrient restriction, in bacteria.

Canonical stress granules are a response to impaired polysome disassembly and are readily reversible. The condensed ring of stress granules around the aggresome is also reversible, although the process is approximately an order of magnitude slower than for dispersed stress granules (Fig. 6c). Hence, the inference is that the highly ordered CCR is a transient response to scarce energy resources and impaired mitochondrial translation, which maintains stress granules and other important structures apart close to the aggresome, ready for dissolution or degradation when energy homeostasis is restored. Meanwhile, the CCR likely helps to suspend many energy-demanding processes, and as it encapsulates the centrosome, mitosis is expected to be the principle restrained activity.

Many methods of inducing protein aggregation result in multiple bodies in the cytoplasm, whereas the inhibitors applied here create a single structure adjacent to the nucleus, akin to those seen in the ALS-FTD disease state. To our knowledge, restricted glycolysis and impaired mitochondrial translation is, the only form of cellular stress to produce stress granules that encircle the aggresome.

The appearance of the TDP-43/FUS rings specifically in the activated microglia of 2DG treated mice is intriguing (Fig. 7), as in ALS, motor neurons are the principal affected cell type, while in FTD it is cortical neurons. One possibility is that the ALS-FTD disease cascade begins with short-lived microglia, and subsequently spreads to motor or cortical neurons. This is all the more credible given that ALS and FTD are among the best examples of spreading proteinopathies (Cicardi et al., 2021) and all cells are phagocytic. Moreover, TDP-43 regulates the phagocytic activity of microglia (Paolicelli et al., 2017; Quek et al., 2022) and stress granule assembly directly affects inflammasome formation and phagocytosis of microglia (Ghosh and Geahlen, 2015; Samir et al., 2019; Wu et al., 2023). Microglia may be particularly vulnerable to restricted glucose utilization, as their glucose uptake increases upon activation (Gimeno-Bayón et al., 2014), and so this might trigger the formation of perinuclear stress granules in the presence of 2DG. Alternatively, microglia may be prone to develop the persistent stress granule rings and inclusions because they clear neuronal TDP-43, as implied by mouse models of ALS where microglial impairment compromises this process (Spiller et al., 2018; Zhang et al., 2020). In the latter case, the microglia abnormality induced by 2DG exposure would mimic a pre-clinical phase of the disease, which would explain the absence of overt neurological dysfunction in the one-year-old mice treated intermittently with 2DG for 8 months.

By defining a unique structure in which many ALS proteins participate, we also offer a solution to the enigma of how mutations in proteins that ordinarily have different locations, such as TDP-43, FUS, VCP, SOD1 and SQSTM1, can contribute to the same disease. Moreover, the profound impact of perturbed energy metabolism on multiple ALS factors suggests restricted nutrient and energy metabolism has a major role in the disease and may in some cases stand at the head of the disease cascade. The idea that disturbed nutrient metabolism can trigger ALS fits well with the disease most often being sporadic, as this could take many different forms, in which genetic background plays only a part. It is known that external factors, such as viral infection and nutrient availability, have marked effects on mitochondria and mitochondrial DNA, and thus energy production (Durigon et al., 2018; Pantic et al., 2021; West et al., 2015). The finding that perturbed energy metabolism can determine the location of a suite of proteins whose dysfunction causes ALS-FTD places restricted glycolysis and mitochondrial protein synthesis center stage in the regulation of protein translocation related to ALS-FTD. As a corollary, targeting mitochondrial translation, respiratory chain capacity or glycolysis, including via modulating nutrient availability, might delay, mitigate or attenuate these diseases.

## METHODS

### Cell culture and treatments

Primary human fibroblasts with or without mutant mitochondrial DNA (m.3243G) were routinely maintained in Dulbecco’s Modified Eagle’s Medium (DMEM) (Gibco) containing 25 mM glucose, 1 mM pyruvate supplemented with 10% fetal bovine serum (FBS, Gibco), 5% penicillin and streptomycin (Gibco), 2 mM GlutaMAX (Gibco), 5% CO_2_, at 37°C. All the cell lines were regularly confirmed free of mycoplasma, using the Venor Gem Classic Mycoplasma PCR Detection Kit (Minerva Biolabs).

The regime for serum-deprived cells was to grow the fibroblasts to 100% confluent and change the medium to DMEM 25 mM glucose, 1 mM pyruvate with 0.1% FBS, which was replaced every 3 days up to day 14. Cells were treated with different compounds and reagents as indicated in the text and figure legends. Chemicals used were 2-Deoxy-D-glucose (2DG) (Sigma Aldrich, Alfa Aesar), Rotenone (Rot) (Glentham life sciences), Carbonyl cyanide 3-chlorophenyl hydrazone (CCCP) (Sigma Aldrich), Chloroquine diphosphate salt (CLQ) (Sigma Aldrich), MG132 (Sigma Aldrich), Epoxomicin (EPO) (Sigma Aldrich), Koningic acid (KA) (Glentham life sciences, Abcam, Santa Cruz), Actinonin (ACT) (Glentham life sciences, Santa Cruz), Chloramphenicol (CAP) (Sigma Aldrich), Cycloheximide (CHX) (Santa Cruz), Dinaciclib (Santa Cruz), Tunicamycin (TN) (Sigma Aldrich), Kifunensine (KIF) (BiosynthCarbosynth), Harringtonine (HARRI) (TargetMol), 1,6-Hexadienol (1,6-HD) (Sigma Aldrich), Nocodazole (NOCO) (Santa Cruz), DBeQ (Santa Cruz), Tubastatin A hydrochloride (TA) (BiosynthCarbosynth), Verdinexor (KPT-335) (TargetMol), TASG (was synthesized in-house according to (Shao et al., 2021)), Sodium arsenite (Sigma Aldrich).

### Immunofluorescence

Fibroblasts were grown on 24-well plates (Corning) with 12 mm #1.5 coverslips (Thermo Fisher Scientific), or on 96-well black microplates (Ibidi), and fixed with 4% paraformaldehyde (Electron Microscopy Sciences) for 20 mins at room temperature (RT). After washing with Phosphate Buffered Saline (DPBS, Gibco), the cells were permeabilized and blocked for 1 h at RT with 10% donkey (GeneTex) or goat serum (Sigma Aldrich) in 0.1% Triton X-100, PBS (PBST). Cells were then incubated with primary antibodies (see Table S1) in PBST overnight (O/N) at 4°C. After three washes for 5 min with PBST, the cells were incubated with the appropriate secondary antibodies (see Table S1) 1:500 dilution in 10% goat/donkey serum in PBST for 1-2 hours at RT. The cells were then washed with PBS, 0.3% Triton X-100, PBS and water, and the coverslips mounted in ProLong® Diamond Antifade Reagent with DAPI (P36962, Thermo Fisher) or in the case of microplates in Mounting medium with DAPI (ibidi). Antibodies were applied according to the manufacturers’ instructions at the dilutions listed in Table S1.

### Image capture and analysis

Fluorescent labeling was observed through a Nikon Eclipse 80i epifluorescence microscope (Figs. 4b, S9D, S11B, S11C), NikonTi Inverted Confocal microscope (Fig. 3d), Nikon Eclipse Ti-5 Inverted epifluorescence microscope (Figs. 1, 2, 4a, S1, S2A, S2B, S4, S5C, S9B) using the NIS elements software or LSM 900 KMAT with AXIO OBSERVER 7 Zeiss confocal microscope (Figs. 3a, 3b, 3e, 3f, 3i, 4c, 5a, 5c, 6, 7, S2C, S2D, S3, S6, S7 S8, S9A, S10, S11A, S12B). Laser power, gain and offset parameters were kept constant for each experiment. The image analysis was performed using the plugins available in Fiji Image J, and all the adjustments were applied equally. The images and illustrations were imported and organized in Adobe Illustrator (Adobe Inc.).

### Fractionation and immunodetection of proteins

Cells were trypsinized and lysed on ice with RIPA buffer (150 mM NaCl, 1.0% TritonX-100, 0.5% Sodium deoxycholate, 0.1% SDS, 50 mM Tris, pH 8.0) 1x Halt™ Protease and Phosphatase Inhibitor Cocktail (Thermo Scientific). After incubating on ice for 40 minutes, the samples were centrifuged for 20 minutes at 15,000 g, 4°C to remove cellular debris. Protein concentration was determined by DC protein assay kit (Biorad). Protein samples were prepared in 1x Laemmli loading buffer (Biorad) with DTT and resolved on 4-15% 4-20% or 12% Mini-PROTEAN® TGX™ Precast Gels (Biorad) run in Tris/Glycine/SDS running buffer (Biorad). After electrophoresis, proteins were transferred to Low-Fluorescence PVDF Transfer Membranes (Thermo Scientific) and blocked in 3% BSA in PBS, 0.1% Tween for 1 h at RT. Membranes were incubated O/N with primary antibodies (see Table S1) in BSA 3% in PBS, 0.1% Tween, at 4°C and, after washing, with the appropriate secondary antibody (see Table 1) for 1 h at RT. Proteins were detected using standard ECLTM Western Blotting Analysis System (GE Healthcare) or SuperSignal®West Dura (Thermo Scientific), and acquired and quantified using an iBright FL1500 Imaging System.

### Nascent protein synthesis

Cells were incubated in methionine-free medium (DMEM without L-methionine and L-cysteine (Cat. no. 21013) supplemented with L-Cysteine dihydrochloride (Alfa Aesar), pyruvate (Gibco), GlutaMAX (Gibco) and 0.1% or 10% FBS, at 37°C and 5% CO_2_ for 30 mins to deplete methionine reserves, 20 mins with or without a cytosolic protein synthesis inhibitor (CHX, 50 μg/mL), and then for 3-4 h with 50 μM Click-iT® HPG (L-Homopropargylglycine) (Invitrogen). For gel electrophoresis analysis, cells were lysed in 1% SDS in 50 mM Tris-HCl, pH 8, or RIPA buffer. 200 μg of total protein in a maximum volume of 50 μL was used for the Click-it reaction between Click-iT® HPG alkyne and Biotin Azide (PEG4 carboxamide-6-Azidohexanyl Biotin) (Invitrogen) was performed using Click-iT™ Protein Reaction Buffer Kit (Invitrogen), and labeled proteins recovered by chloroform/methanol extraction, and resuspended in RIPA buffer according to the manufacturer’s protocol. To separate HPG labeled proteins via SDS gel electrophoresis (5 μg protein/lane), we found it was essential to include the reducing agent dithiothreitol, 100 mM DTT in 4 x loading buffer, even though the manufacturer expressly discourages this. After transfer to PVDF membrane, biotin azide (with HPG alkyne) was detected by Streptavidin Alexa Fluor™ 647 conjugate (Invitrogen) and acquired by iBright FL1500 Imaging. For cell imaging, cells plated on 96-well microplates (Ibidi) were fixed with 4% paraformaldehyde (Electron Microscopy Sciences) for 20 mins at RT, and the Click-it reaction between Click-iT® HPG alkyne (present in protein and Alexa Fluor™ 555 Azide, Triethylammonium Salt (Invitrogen) was applied using the Click-iT™ Cell Reaction Buffer Kit (Invitrogen). Parallel immunostaining (e.g., of the mitochondria network with anti-TOMM20 antibody) was performed as normal.

### Statistical Analysis

Normal distribution was determined by the Shapiro-Wilk test. For normally distributed data single comparisons were tested using the independent Student’s t-test. Non-parametric data were analyzed by the Mann-Whitney U-test. Statistical significance was set at \**p*≤ 0.05, \*\**p*≤ 0.01, \*\*\**p*≤ 0.001, or <0.0001. The data are presented as means ± SD. Statistical analyses were performed using GraphPad Prism 8.0.

### ^35^S-methionine in-cell assay of mitochondrial protein synthesis

Mitochondrial translation products were labeled using ^35^S-methionine as described previously (Durigon et al., 2018). Briefly, fibroblasts were washed twice with methionine-free medium and incubated in the same medium for 10 min at 37°C. 100 μg/ml Emetine dihydrochloride (Sigma) was added to inhibit cytosolic translation, before pulse-labeling with 100 μCi [^35^S]-methionine for 60 min. Cells were chased for 10 min at 37°C in regular DMEM with 10% FBS, washed three times with PBS, and collected. Labeled cells were lysed in a solution of PBS, 0.1% n-dodecyl-D-maltoside (DDM), 1% SDS, 50 units of benzonase (Novagen), and 1:50 (v/v) Roche protease inhibitor cocktail. Protein concentration was measured by Lowry assay, and 20 μg of protein was separated by 12% SDS–PAGE. Gels were dried and exposed to phosphor plates, and the signal detected via a PhosporImager.

### VCP silencing

For RNAi experiments, primary human fibroblasts were transfected with siRNA Reagent System (Santa Cruz) and VCP siRNA (h) (sc-37187) (Santa Cruz) or Control siRNA-A (siCA) (sc-37007) (Santa Cruz), the latter as a negative control. Immunostaining was performed 96 h after transfection.

### Live cell imaging

For Golgi live imaging, human primary fibroblasts were transfected with CellLight™ Golgi-GFP, BacMam 2.0 (Invitrogen) for 16 h, and time-lapse imaging was performed in FluoroBrite™ DMEM (Gibco) (with or without Hoechst 1 μg/mL) in AXIO OBSERVER 7 Zeiss confocal microscope PECON XL incubator, 5% CO_2_, at 37°C. For G3BP1 live imaging a GFP-G3BP1 fusion protein containing plasmid (pEGFP-C1-G3BP1-WT, Addgene plasmid # 135997) was transfected with Lipofectamine™ 3000 Transfection Reagent (Invitrogen). 72 h after the transfection, GFP-G3BP1 signal was visualized in FluoroBrite™ DMEM (Gibco) in the same microscope. Subsequently, the cells were fixed and immunostained with anti-G3BP1 antibody.

### Proteasome activity in live cells

Proteasome-Glo™ Chymotrypsin-Like, Trypsin-Like and Caspase-Like Cell-Based Assay (Promega) and GloMax® Discover microplate reader (Promega) was used to measure the chymotrypsin-like, trypsin-like or caspase-like protease activity associated with the proteasome complex in cultured fibroblasts. Proliferative cells were seeded in 384-well plate 48 h before the assay was performed, whereas quiescent cells were obtained as described above in the same 384-well plate having been seeded 13 days earlier. All cells were treated with different treatments 24 h before the assay. All conditions were done in triplicates. Positive proteasome inhibitors MG132 and EPO were used. After the assay cells were fixed with 4% PFA (E.M.) 10 min RT and nucleus were stained with Ibidi mounting medium with DAPI (Ibidi). DAPI positive cells were counted by microscope imaging and processing by ImageJ software. Luminescence data of each proteasome activity assay was normalized with respect to cell number, and the luminescence of untreated cells of each cell line.

### Transmission electron microscopy

Cells in a Nunc® Lab-Tek® 8-well chamber slide (Merck) were washed with Phosphate Buffer (PB) 0.1 M pH 7.2 (NaH_2_PO_4_·H_2_O and Na_2_HPO_4_) and fixed with 3% glutaraldehyde (Electron Microscopy Sciences) for 10 min at 37°C and 2 h at RT. After five further washes of 5 min, the chamber slides were dried and the samples cut and processed (Centro de Investigación Principe Felipe, Valencia). Electron microscope images were taken with a 200 kV high-resolution TECNAI G2 20 TWIN transmission electron microscope (Sgiker, University of the Basque Country).

### Mice treatment and tissue analysis

Animal handling was conducted following the European Council Directive 2010/63/UE. The Ethical Committee and implied regulatory offices and local authorities approved all experimental procedures at CIC biomaGUNE (authorization number: PRO-AE-SS-185). C57BL/6 mice of three months of age were purchased from Charles River laboratories and were fed a standard chow, and normal unadulterated drinking water (control untreated mice), or drinking water with 0.4% (w/v) 2DG three consecutive days per week, which based on liquid intake equated to ∼400 mg/Kg body weight per day, on the days when 2DG was present in the drinking water. After 8 months, mice were sacrificed 24 h after 2DG was withdrawn for the final time. Tissue samples were fixed in 4% paraformaldehyde for 24 h and embedded in paraffin. Coronal brain sections of 4 µM thickness were cut and mounted on coverslips for staining. Tissue sections were stored at room temperature. Samples were deparaffinized by a standard procedure using xylol (15 minutes) and a graded series of ethanol washes (5 minutes each). After incubation with PBS-Tween 0.1 % (PBST) samples were submerged with the antigen retrieval Sodium Citrate Buffer pH 6 for 30 minutes, at 95°C. Sections were then rinsed in PBST (2 x 3 minutes) and blocked with a solution containing 10% Goat Serum in PBST for 1 h. Next, sections were incubated overnight at 4°C with the appropriate primary antibody. Next day samples were washed with PBST (3 x 5 minutes) and incubated for 1 h with the secondary antibody and DAPI (5 µg/mL). Samples were mounted with mounting medium ProLong™ Diamond AntifadeMountan with DAPI (Life technologies) and covered with a #1.5 coverslip. Antibodies were applied according to the manufacturers’ instructions at the dilutions listed in Table S1.

## ACKNOWLEDGEMENTS

UFP, ALA and AE were supported by predoctoral fellowships from the Basque Government (PRE_2018_1_0253, PRE_2019_1_0184 and PRE_2020_1_0119) and MMO was the recipient of a predoctoral fellowship from the University of the Basque Country (PIF18/317). MMO was also partially supported by the Basque Government’s IKUR strategy Neurodegenprot project. IJH was supported by funding from the Spanish Ministry of Health (ISCIII: PI17-00380; PI20/00096) and the Basque Government Department of Health (grants 2021111070; 2022333050; 2018111043; 2018222031). AS receives support from Brain Research UK. SAM is supported by the Gipuzkoa Fellow of Talent Attraction and Retention from Diputación Foral de Gipuzkoa (2019-FELL-000010-01, 2020-FELL-000016-02-01, 2021-FELL-000013-02-01). FJGB was supported by Roche Stop Fuga de cerebros (BIO19/ROCHE/017/BD). JRC is supported by grants from the Ministry of Science, Innovation and Universities (SAF2017-84494-C2-R), the Gobierno Vasco, Dpt. Industry, Innovation, Commerce & Tourism under the ELKARTEK Program (KK-2019/bmG19). JRC also received funding from the BBVA Foundation (Ayudas a Equipos de investigación científica Biomedicina 2018) and from La Caixa Foundation (Health Research Call 2020: HR20-00075) and the Maria de Maeztu Units of Excellence Programme (MDM20170720).

## Ethics statements

The human cell lines used in this study were obtained after having been approved by the appropriate institutional and regional ethics committee and the experiments and data handling have been performed in accordance with the ethical standards. The animal part of the study was approved by the Ethical Committee of CIC biomaGUNE and local authorities of Gipuzkoa (PRO-AE-SS-185). Animal maintenance and handling was conducted in accordance with the European Council Directive 2010/63/UE.

